# Genome editing enables reverse genetics of multicellular development in the choanoflagellate *Salpingoeca rosetta*

**DOI:** 10.1101/2020.02.18.948406

**Authors:** David S. Booth, Nicole King

**Affiliations:** Howard Hughes Medical Institute and Department of Molecular and Cell Biology, University of California, Berkeley, CA 94720; Department of Biochemistry and Biophysics, University of California, San Francisco, CA 94158

## Abstract

In a previous study, we established a forward genetic screen to identify genes required for multicellular development in the choanoflagellate, *Salpingoeca rosetta* (Levin et al., 2014). Yet, the paucity of reverse genetic tools for choanoflagellates has hampered direct tests of gene function and impeded the establishment of choanoflagellates as a model for reconstructing the origin of their closest living relatives, the animals. Here we establish CRISPR/Cas9-mediated genome editing in *S. rosetta* by engineering a selectable marker to enrich for edited cells. We then use genome editing to disrupt the coding sequence of a *S. rosetta* C-type lectin gene, *rosetteless*, and thereby demonstrate its necessity for multicellular rosette development. This work advances *S. rosetta* as a model system in which to investigate how genes identified from genetic screens and genomic surveys function in choanoflagellates and evolved as critical regulators of animal biology.

## Introduction

As the sister group of animals (Fig. 1A), choanoflagellates have great potential for revealing the origins of animal development and the cell biology of multicellularity (Lang et al., 2002; Burger et al., 2003; Carr et al., 2008; Ruiz-Trillo et al., 2008; Grau-Bové et al., 2017). Comparative genomic studies have demonstrated that choanoflagellates express genes that are necessary for animal development (King et al., 2008; Fairclough et al., 2013; Richter et al., 2018), including genes for intercellular adhesion (e.g. cadherins: Abedin and King, 2008; Nichols et al., 2012), signaling (e.g. receptor tyrosine kinases and CamKII: Manning et al., 2008; Pincus et al., 2008; Bhattacharyya et al., 2016; Amacher et al., 2018), and cellular differentiation (e.g. myc, STAT, and p53: Young et al., 2011; Mendoza et al., 2013). Moreover, choanoflagellates and animals are the only clades that have cells with a collar complex (Leadbeater, 2015; Brunet and King, 2017), a unique cellular module in which a collar (*choano* in Greek) of actin-filled microvilli surrounds an apical flagellum (Fig. 1B; Sebé-Pedrós et al., 2013; Peña et al., 2016; Colgren and Nichols, 2019). Together, these observations have motivated the development of choanoflagellates as models for researching the function and evolution of core developmental regulators (King, 2004; Hoffmeyer and Burkhardt, 2016; Sebé-Pedrós et al., 2017; Brunet and King, 2017).

**Fig. 1:**
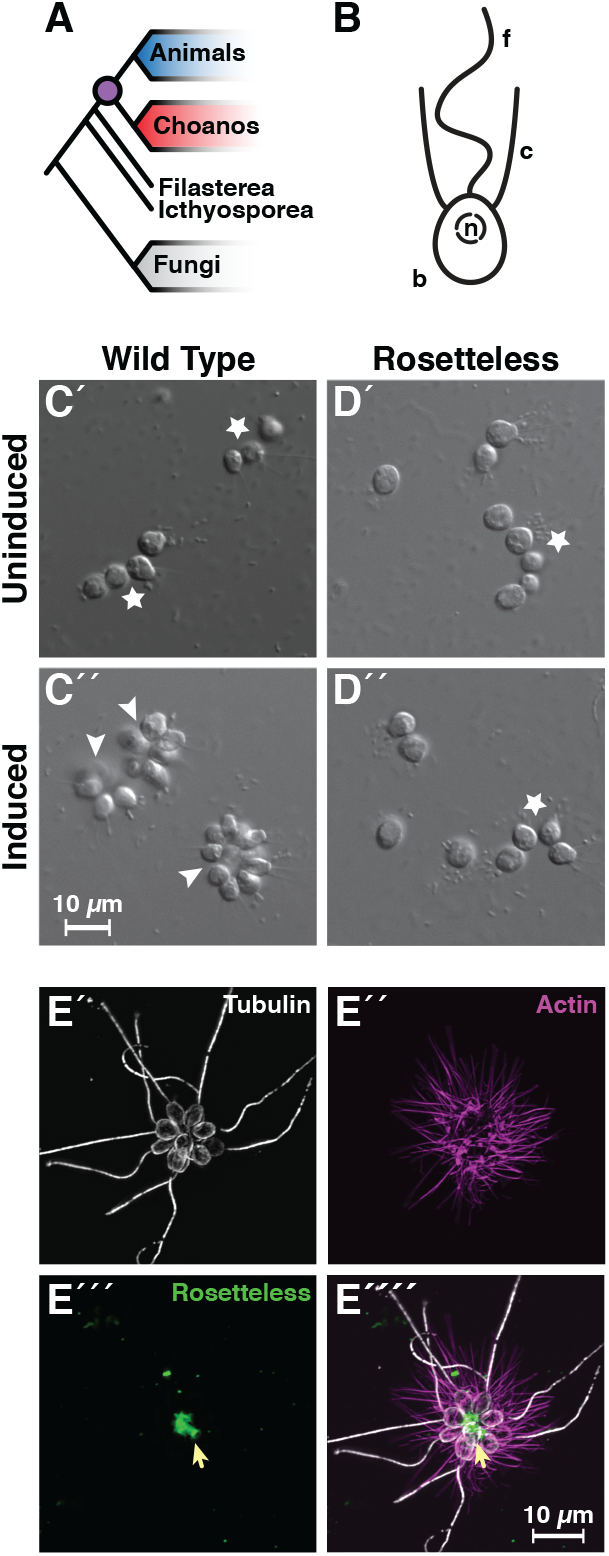
Genome editing in *S. rosetta* establishes reverse genetics in choanoflagellates, the closest living relatives of animals. (A) Choanoflagellates (blue) are the closest living relatives of animals (red) and last shared a common ancestor (purple) ∼800 million years ago (Parfrey et al., 2011). (B) The collar complex, an apical flagellum (f) surrounded by a collar (c) of actin-filled microvilli, typifies choanoflagellates and is uniquely shared between choanoflagellates and animals (Brunet and King, 2017). (C) Wild-type *S. rosetta* forms multicellular rosette colonies in response to rosette inducing factors (RIFs) secreted by environmental bacteria. In the absence of RIFs (C’), *S. rosetta* grows as single cells or as a linear chain of cells (star). Upon the addition of RIFs (C”; Alegado et al., 2012; Woznica et al., 2016), *S. rosetta* develops into spheroidal, multicellular rosettes (arrowhead) through serial cell divisions (Fairclough et al., 2010). (D) The *rosetteless* C-type lectin gene is necessary for rosette development. A mutation in *rosetteless* allows normal cell growth as single cells and linear chains in the absence of RIFs (D’) but prevents rosette development in the presence of RIFs (D”; Levin et al., 2014). (E) Wild-type *S. rosetta* secretes Rosetteless protein from the basal ends of cells into the interior of rosettes. Shown is a representative rosette stained with an antibody to alpha-tubulin to mark cortical microtubules and the apical flagellum of each cell (E’, grey) phalloidin to mark actin-filled microvilli (E”, magenta), and an antibody to Rosetteless protein (E’’’, green). A merge of alpha-tubulin, phalloidin, and Rosetteless staining shows that Rosetteless protein localizes to the interior of rosettes (arrow) where cells meet at their basal ends (E’’’’; Levin et al., 2014).

The choanoflagellate *Salpingocea rosetta* has received the greatest investment in tool development (Hoffmeyer and Burkhardt, 2016). Its 55.44 megabase genome encodes ∼11,629 genes, some of which are homologs of integral regulators for animal development (Fairclough et al., 2013). Moreover, the life history of *S. rosetta* provides a rich biological context for investigating the functions of intriguing genes (King et al., 2003; Fairclough et al., 2010; Dayel et al., 2011; Levin and King, 2013; Woznica et al., 2017). For example, *S. rosetta* develops into multicellular, spheroidal colonies called rosettes through serial cell divisions from a single founding cell (Fairclough et al., 2010; Laundon et al., 2019; Larson et al., 2020), a process induced by environmental bacteria that can also serve as a food source (Fig. 1C; Alegado et al., 2012; Woznica et al., 2016). Thus, rosette development can provide a phylogenetically relevant model for discovering genes that mediate multicellular development and bacterial recognition in choanoflagellates and animals.

A forward genetic screen was established to hunt for mutants that were unable to develop into rosettes and resulted in the identification of genes required for rosette development (Levin et al., 2014; Wetzel et al., 2018). The first of these (Levin et al., 2014), *rosetteless* encodes a C-type lectin protein that localizes to the interior of rosettes (Fig. 1D-E). As C-type lectins are important for mediating intercellular adhesion in animals (Drickamer and Fadden, 2002; Cummings and McEver, 2015), this discovery highlighted the conserved role of an adhesion protein family for animal and choanoflagellate development. However, the screen also underscored the necessity for targeted genetics in *S. rosetta*. Because of inefficient mutagenesis in *S. rosetta*, forward genetics has been laborious: out of 37,269 clones screened, only 16 rosette-defect mutants were isolated and only three of these have been mapped to genes (Levin et al., 2014; Wetzel et al., 2018). Establishing genome editing would accelerate direct testing of gene candidates identified through forward genetic screens, differential gene expression, and/or genomic comparisons.

Therefore, for the present study, we sought to establish CRISPR/Cas9 genome editing in *S. rosetta*. Cas9-mediated genome editing (Jinek et al., 2012, 2013; Cong et al., 2013) has been crucial for advancing genetics in emerging models (Gilles and Averof, 2014; Harrison et al., 2014; Momose and Concordet, 2016). Depending on the DNA repair pathways expressed in a given organism (Yeh et al., 2019), the delivery of the Cas9 endonuclease bound to a programmable guide RNA (gRNA) can direct DNA cleavage at a target site to introduce mutations from co-delivered DNA templates or from untemplated repair errors that cause insertions or deletions (Rouet et al., 1994; Choulika et al., 1995; Bibikova et al., 2001; Jinek et al., 2013; Cong et al., 2013). While the delivery of macromolecules into choanoflagellate cells has been a longstanding barrier for establishing reverse genetic tools, we recently established a robust method to transfect *S. rosetta* with DNA plasmids for expressing transgenes (Booth et al., 2018), which allowed us to perform genetic complementation (Wetzel et al., 2018). Despite having established a method for gene delivery in *S. rosetta*, the lack of knowledge about DNA repair mechanisms in choanoflagellates and low-transfection efficiency (∼1%) presented challenges for establishing genome editing, particularly without a proven selectable marker to enrich for editing events.

Here we report a robust method for genome editing to perform reverse genetics in *S. rosetta*. First, we engineered a selectable marker for cycloheximide resistance as an initial demonstration of genome editing with CRISPR/Cas9 (Fig. 2). We then inserted a foreign sequence into *rosetteless* that eliminates its function, confirming the importance of this gene for multicellular rosette development (Fig. 3). Finally, we found that, even in the absence of selection, *S. rosetta* preferentially uses DNA templates to repair double-stranded breaks (Fig. 4). This work establishes genome editing in *S. rosetta* and provides a path for testing the function of choanoflagellate genes that are implicated in the early evolution of animals.

**Fig. 2:**
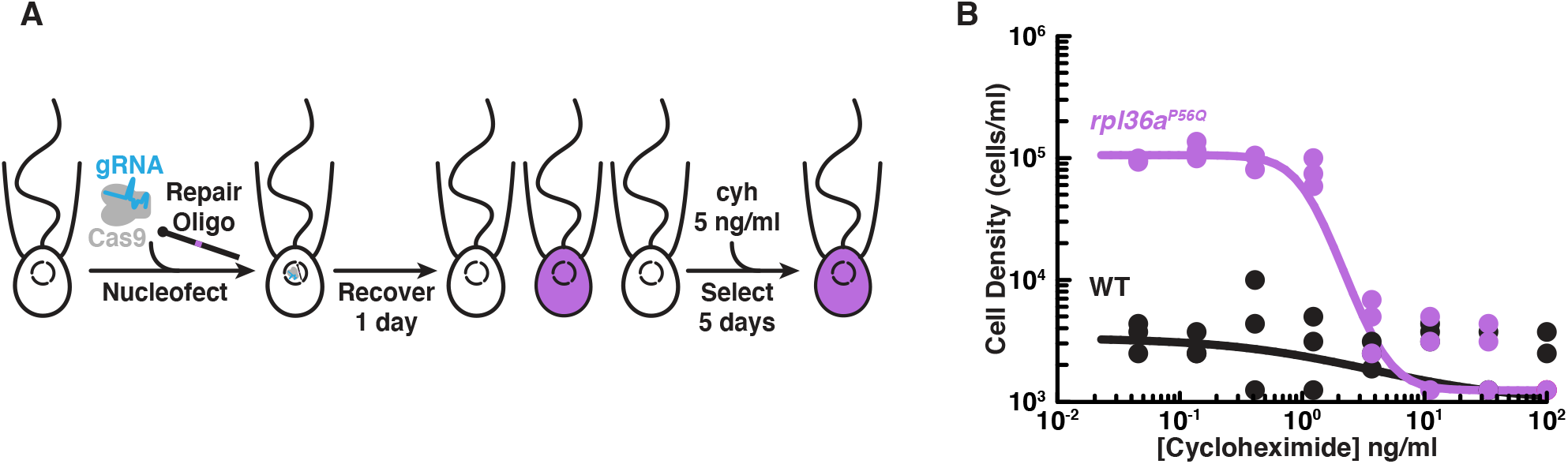
Engineered cycloheximide resistance in *S. rosetta* provides a proof-of-principle for Cas9-mediated genome editing. (A) Schematic of Cas9-mediated genome editing to engineer cycloheximide resistance in *S. rosetta*. Nucleofection was used to deliver *Sp*Cas9 (gray) bound to gRNA (cyan), which together form the *Sp*Cas9 RNP, and repair oligonucleotides (Repair Oligo; Fig. S2) to engineer cycloheximide resistance. After recovering cells for one day, successfully edited cells were selected by growth in media supplemented with cycloheximide (cyh), which inhibits the growth of wild-type cells (Fig. S1) and selects for cycloheximide-resistant cells (purple). (B) A designer cycloheximide-resistant allele (Fig. S2) allows cell proliferation in the presence of cycloheximide. Wild-type (WT, black dots and line) and *rpl36a^P56Q^* (purple dots and line) strains were placed into media supplemented with a range of cycloheximide concentrations (x-axis) at a cell density of 10^4^ cells/ml and then were grown for two days. *rpl36a^P56Q^* grew to higher cell densities than the wild-type strain at cycloheximide concentrations <10 ng/ml. At higher concentrations, cycloheximide inhibited growth of both strains. The dots show cell densities from three independent replicates. The lines show the average from independently fitting a dose inhibition curve to the cell densities from three independent experiments.

**Fig. 3:**
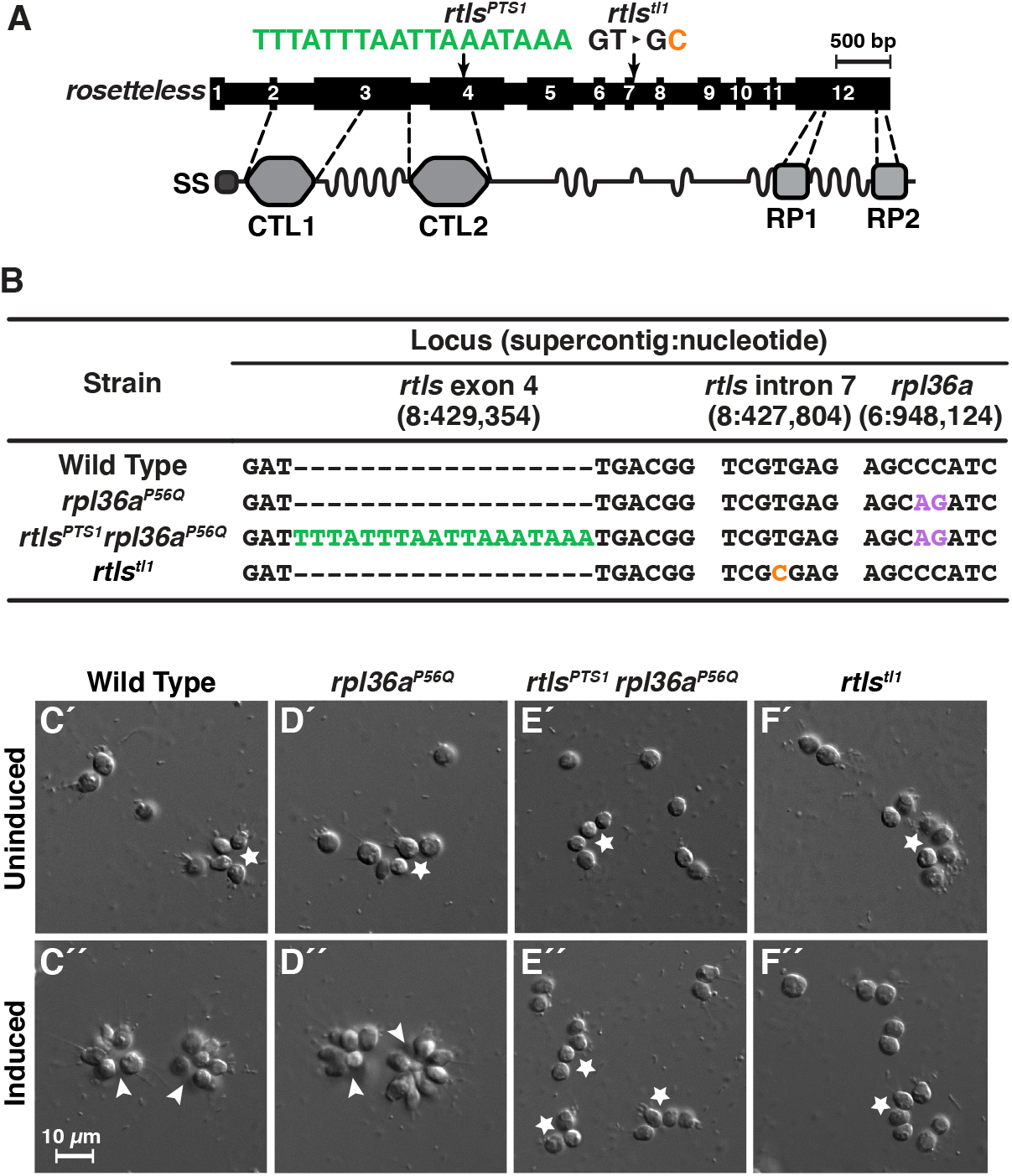
Genome editing of *rosetteless* enables targeted disruption of multicellular development in *S. rosetta*. (A) An engineered mutation in *rosetteless* introduces a premature termination sequence (PTS) to knockout the expression of *rosetteless*. The *rosetteless* gene (exons shown as numbered black boxes, connected by introns) encodes a secreted protein (SS denotes the signal sequence for secretion) with two C-type lectin domains (CTL1 and CTL2) and two carboxy-terminal repeats (RP1 and RP2). A forward genetic screen (Levin et al., 2014) identified a mutation, *rtls^tl1^*, in which a T to C transition in the seventh intron disrupts splicing and knocks out *rosetteless* expression. To increase the likelihood of disrupting *rosetteless* function with genome editing, we designed the *rtls^PTS1^* mutation that introduces a PTS (green), with a poly-adenylation sequence and stop codons in each reading frame, into the fourth exon of the gene. (B) The genotypes of strains established from genome-editing confirm that *rosetteless* and *rpl36a* incorporated the designed mutations. To enrich for genome-edited cells, *Sp*Cas9 RNPs and repair templates for introducing *rpl36a^P56Q^* (Fig. S2) and *rtls^PTS1^* were simultaneously delivered into *S. rosetta*. Afterward, cycloheximide resistant cells were clonally isolated and screened for cells that did not develop into rosettes in the presence of RIFs. The genotypes of *rtls^PTS1^ rpl36a^P56Q^*, and *rpl36^P56Q^* confirmed that strains established from genome editing had the *rpl36^P56Q^* allele and the strain with the *rosetteless* phenotype also had the *rtls^PTS1^* allele. In addition, the wild-type and genome edited strains lacked the T to C transition in the 5’-splice site of intron 7 that defined the *rtls^tl1^* allele. (C-F) Phenotypes of genome-edited strains correspond to their respective genotypes. In the absence of RIFs, all strains (C’, D’, E’, and F’) grew as chains (stars) or single cells. Upon the addition of RIFs, the wild-type (C”) and *rpl36^P56Q^* strains (D”) formed rosettes and expressed the Rosetteless protein (Fig. S3). In contrast, *rtls^PTS1^ rpl36^P56Q^* (E”) and *rtls^tl1^* (F”) did not form rosettes and did not express Rosetteless (Fig. S3).

**Fig. 4:**
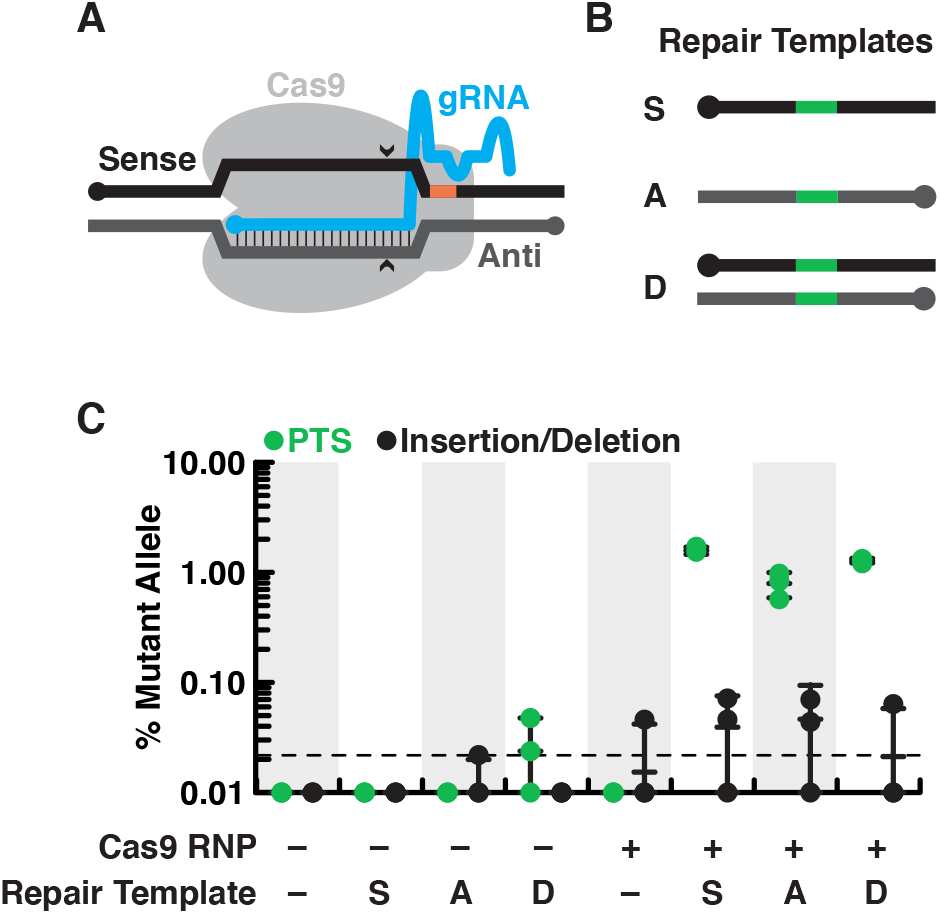
*S. rosetta* preferentially repairs *Sp*Cas9 cleavage with DNA templates. (A) Schematic of a gRNA targeting *Sp*Cas9 to a genomic locus of interest. A gRNA (cyan, knobs indicate 5’ ends) that encodes a 20 nt targeting sequence from the sense strand of a genomic locus (black) hybridizes with the antisense strand (dark gray). *Sp*Cas9 (light gray) introduces a double-stranded break at the genomic locus (carets), 3 bp upstream of a protospacer adjacent motif (PAM, orange). (B) We designed a panel of repair oligonucleotides to test the preferred substrates for repairing double-stranded breaks introduced by *Sp*Cas9 at *rosetteles* exon 4. Oligonucleotide repair templates containing the PTS sequence (green) were delivered as single-stranded DNA in the sense (S) or anti-sense (A) orientations and as a double-stranded template (D) to test which most efficiently templated DNA repair at the *Sp*Cas9 cleavage site. (C) *Sp*Cas9 stimulated the repair from DNA templates. Repair templates with a PTS (from panel B) were delivered in the presence and absence of *Sp*Cas9 (+/–). A ∼450 bp fragment surrounding the *rtls^PTS1^* cleavage site was amplified from cells that had been transfected the previous day to prepare deep sequencing libraries for quantifying the frequency of PTS insertions (green) or insertions/deletions from error prone editing (black). Each experiment was performed three independent times (dots; mean and standard deviations are shown with lines). The dotted line indicates the limit of detection of the sequencing, based on a 6 base, randomized barcode. Upon transfection with the *Sp*Cas9 RNP, 10x more mutations from repair templates (1-2%, green dots) were detected than untemplated insertions or deletions (black dots). In the absence of *Sp*Cas9, mutations generated from a double-stranded template, but not single-stranded templates, were rarely (<0.1%) and unreliably (2 of 3 trials) found.

## Results

### A marker to select for cycloheximide resistance facilitates genome editing in *S. rosetta*

Our initial attempts to target rosetteless for genome editing in *S. rosetta* were either unsuccessful or resulted in editing events that were below the limits of detection. Therefore, suspecting that genome editing in *S. rosetta* might prove to be challenging to establish, we first aimed to introduce a mutation in an endogenous gene that would confer antibiotic resistance and allow selection for rare genome editing events.

In *Chlamydomonas* (Stevens et al., 2001) and Fungi (Kawai et al., 1992; Dehoux et al., 1993; Kondo et al., 1995; Kim et al., 1998), specific mutations in the ribosomal protein gene *rpl36a* confer resistance to the antibiotic cycloheximide by disrupting cycloheximide binding to the large subunit of eukaryotic ribosomes (Stöcklein and Piepersberg, 1980; Schneider-Poetsch et al., 2010; Garreau de Loubresse et al., 2014). After finding that *S. rosetta* cell proliferation was inhibited by cycloheximide (Fig. S1A), we sought to establish a cycloheximide-resistant strain through genome editing. By combining prior genetic findings (Bae et al., 2018) with our own structural modeling (Fig. S1B) and bioinformatic analyses (Fig. S1C) of the *S. rosetta rpl36a* homolog (PTSG_02763), we predicted that converting the 56^th^ codon of *rpl36a* from a proline to a glutamine codon (*rpl36a^P56Q^*) would render *S. rosetta* resistant to cycloheximide. Insertion or deletion mutations that could arise as errors from repairing the double-stranded break without a template would likely kill cells by disrupting the essential function of *rpl36a* for protein synthesis (Bae et al., 2018).

To edit the *rpl36a* gene in *S. rosetta*, we first designed a gRNA with a 20 nt sequence from *rpl36a* to direct Cas9 from *Streptomyces pyogenes* (*Sp*Cas9) to cut at *S. rosetta* supercontig 6: 948,122 nt (Fairclough et al., 2013). Then we made a DNA repair template as a single-stranded DNA oligonucleotide with a sequence encoding the Pro56Gln mutation and 200 bases of flanking homologous sequence from *rpl36a* centered on the cleavage site (Fig. S2A-B). To deliver the *Sp*Cas9/gRNA ribonucleoprotein complex (*Sp*Cas9 RNP) and the repair template encoding the Pro56Gln mutation into *S. rosetta* cells, we used a nucleofection protocol adapted from our recently developed method for transfecting *S. rosetta* (Fig. 2A; Booth et al., 2018). We favored delivering the *Sp*Cas9 RNP rather than expressing *Sp*Cas9 and gRNAs from plasmids, as overexpressing *Sp*Cas9 can be cytotoxic for other organisms (Jacobs et al., 2014; Jiang et al., 2014; Shin et al., 2016; Foster et al., 2018) and RNA polymerase III promoters for driving gRNA expression have not yet been characterized in *S. rosetta*. After growing transfected cells in the presence of cycloheximide for five days, Sanger sequencing of PCR-amplified *rpl36a* showed that *rpl36a^P56Q^* was the major allele in the population (Fig. S2, compare B and C). Sequencing a clonal strain established from this population confirmed the *rpl36a^P56Q^* genotype (Fig. S2D), and growth assays showed that the *rpl36a^P56Q^* strain proliferated better than the wild-type strain in increasing concentrations of cycloheximide (Fig. 2B).

The ability to engineer cycloheximide resistance additionally offered a simple assay to optimize essential parameters for genome editing in *S. rosetta*. Therefore, we tested how varying delivery conditions would impact the frequency of template-mediated mutagenesis and, ultimately, the cell density and consensus genotype of a cell population after genome editing and cycloheximide treatment (Fig. S2E-H). Through this optimization process (Fig. S2), we found that efficient genome editing required transfection with at least 20 pmol of *Sp*Cas9 RNP and more than 200 nmol of a single-stranded DNA repair template that had 50 bases of homology flanking a designed mutation. Henceforth, these parameters established baseline conditions for designing and executing genome editing experiments.

### Targeted disruption of *rosetteless* demonstrates its essentiality for multicellular rosette development

We next sought to use genome editing as a general tool for reverse genetics in choanoflagellates. To this end, we targeted *rosetteless* (*rtls*), one of only three genes known to be required for rosette development in *S. rosetta* (Levin et al., 2014; Wetzel et al., 2018). A prior forward genetic screen linked the first rosette defect mutant to an allele, *rtls^tl1^*, in which a T to C transition in the 5’-splice site of intron 7 (*S. rosetta* supercontig 8: 427,804 nt; Fairclough et al., 2013) was associated with the disruption of *rtls* expression and rosette development (Fig. 3A and 1C-E; Levin et al., 2014). We therefore sought to generate a new *rtls* knockout allele, whose phenotype we predicted would be loss of rosette development.

To increase the likelihood of generating a *rtls* knockout through genome editing, we aimed to introduce sequences that would prematurely terminate transcription and translation near the 5’ end of the gene. First, we designed a gRNA that would target *Sp*Cas9 to the 5’ end of *rtls*. Next, we designed a general-purpose premature termination sequence (PTS), an 18-base, palindromic sequence (5’-TTTATTTAATTAAATAAA-3’) that encodes polyadenylation sequences and stop codons on both strands and in each possible reading frame. This sequence should prematurely terminate transcription and translation to either create a gene truncation or fully knockout target gene expression. We then designed a DNA oligonucleotide repair template in which the PTS was inserted into 100 bp of *rtls* sequence centered around the *Sp*Cas9 cleavage site (supercontig 8: 429,354 nt).

The low efficiency of transfection (∼1%; Booth et al., 2018), the inability to select for cells with the Rosetteless phenotype, and the unknown but potentially low efficiency of genome editing meant that it might be difficult to recover cells in which *rosetteless* had been edited. To overcome this challenge, we sought to simultaneously edit *rosetteless* and *rpl36a* by transfecting cells with RNPs complexed with gRNAs and DNA repair templates for both knocking out *rosetteless* and engineering cycloheximide resistance. In other organisms (Arribere et al., 2014; Kim et al., 2014; Ward, 2015; Foster et al., 2018), this approach has allowed for co-selection by using a selectable marker to improve the recovery of cells that contain a second mutation in a different locus. In *S. rosetta*, we found that 10.4-16.5% of cycloheximide resistant cells contained the *rtls^PTS1^* allele when *rosetteless* and *rpl36a* were co-edited (Fig. S3A).

By first selecting for cycloheximide resistance and then performing clonal isolation by limiting dilution, we were able to isolate multiple clonal lines that were resistant to cycloheximide. We focused on one strain that correctly formed rosettes in response to bacterial rosette inducing factors (RIFs; Fig. 3D; Alegado et al., 2012; Woznica et al., 2016) and two cycloheximide-resistant strains that failed to form rosettes in the presence of RIFs (representative strain shown in Fig. 3E). Genotyping of these strains at exon 4 of *rosetteless* and at *rpl36a* (Fig. 3B) showed that: (1) all three cycloheximide resistant strains established from the same genome-edited population had the cycloheximide resistance allele, (2) the strains that developed into rosettes only had the cycloheximide resistant allele, *rpl36a^P56Q^*, and (3) the two strains that did not develop into rosettes also had the PTS in *rosetteless* exon 4, meaning their genotype was *rtls^PTS1^ rpl36a^P56Q^* (Fig. 3B). For comparison, we also genotyped wild-type, *rpl36a^P56Q^*, *rtls^PTS1^ rpl36a^P56Q^*, and *rtls^tl1^* strains at intron 7 of *rosetteless*, where the *rtls^tl1^* mutation was mapped, to further underscore that *rtls^PTS1^* is an independent mutation that prevents the development of rosettes (Fig. 3B).

To further validate the genotype-to-phenotype relationship of the *rosetteless* knockouts (Fig. 3C-F), we analyzed the percentage of cells that developed into rosettes (Fig. S3B), the localization of the Rosetteless protein (Fig. S3C-F), and the rates of proliferation (Fig. S4) in the wild-type, *rpl36a^P56Q^*, *rtls^PTS1^ rpl36a^P56Q^*, and *rtls^tl1^* strains of *S. rosetta*. In each of these assays, the *rtls^PTS1^ rpl36a^P56Q^* strains exhibited the same phenotype as *rtls^tl1^* (Fig. 3, compare E to F): no cells developed into rosettes (Fig. S3B), an anti-Rosetteless antibody did not detect Rosetteless protein at the basal end of cells (Fig. S3, compare E-F to C), and the mutant and wild-type strains proliferated comparably well (Fig. S4, C-E). Furthermore, *rpl36a^P56Q^* developed into wild-type rosettes (Fig. 3, compare D to C and Fig. S4A), localized Rosetteless protein to the basal end of cells (Fig. S3, compare D to B), and proliferated as rapidly as the wild-type strain (Fig. S4A-B, E), demonstrating that the act of genome editing alone does not yield non-specific defects in rosette development. Our ability to engineer a new *rosetteless* allele, *rtls^PTS1^*, that mimics the rosette-defect phenotype of *rtls^tl1^* demonstrates the potential of genome editing as a general tool for generating targeted gene knockouts in choanoflagellates.

### S. rosetta preferentially repairs double-stranded breaks with DNA templates for genome editing

Thus far, we had only detected mutations from repair templates with homology arms spanning both sides of the double-strand break (Figs. 2, 3, and S2). However, selecting for cycloheximide resistance may have favored those repair outcomes, as insertion and deletion mutations arising from untemplated repair are likely to be deleterious for the function of *rpl36a*. Therefore, to investigate the frequency of template-mediated repair in the absence of selection, we sought to edit *rosetteless*, which is not required for viability (Fig S5).

As prior work has shown that editing outcomes in different cell types (Harrison et al., 2014; Yeh et al., 2019) can depend on the length and orientation (anti-sense or sense) of homology arms (Lin et al., 2014; Richardson et al., 2016; Paix et al., 2017; Li et al., 2019; Okamoto et al., 2019) and chemical modifications of DNA repair templates (Renaud et al., 2016; Yu et al., 2019), we designed a panel of diverse double- and single-stranded DNA repair templates (Fig. 4B and Fig. S5). The double-stranded templates contained phosphorylation or phosphorothioate bonds at their 5’ ends (Fig. S5A); whereas, the single-stranded templates varied in their orientation and presence of 5’ or 3’ homology arms (Fig. S5B). We transfected cells with these repair templates with or without the *Sp*Cas9 RNP. After the cells recovered for one day, we amplified a ∼450 bp fragment around the *Sp*Cas9 cut site for deep sequencing (Lin et al., 2014) and quantified the frequency and type of mutation after genome editing.

We found that *S. rosetta* could use a variety of templates with homology arms spanning both sides of the *Sp*Cas9 cleavage site to repair DNA, each time with a frequency of ∼1-2% (Fig. 4C and Fig. S5C). In the presence of the *Sp*Cas9 RNP and a repair template, we detected edits that were incorporated from DNA templates 10x more frequently than insertion and deletion mutations, which occurred at a frequency of <0.1% (Fig. 4C, see Fig. S5 D-E for the types of insertions and deletions). Notably, the total frequency of genome editing (∼1-2 %) is on the same order as transfection efficiency (∼1%; Booth et al., 2018), suggesting that the delivery of the *Sp*Cas9 RNP and repair templates is the biggest factor limiting genome editing efficiency. Although *Sp*Cas9 was essential for efficient mutagenesis with all of the repair templates, we did observe the incorporation of a double-stranded repair template at a frequency of ∼0.02% in the absence of *Sp*Cas9. In the end, our optimization efforts revealed that the initial template design (Fig. 2–3 and Fig. S2), a single-stranded template in the sense orientation with homology arms spanning both sides of the *Sp*Cas9 cleavage site, was most efficient for genome editing.

## Discussion

The establishment of Cas9-mediated genome editing advances *S. rosetta* as a model for illuminating the evolution of development in choanoflagellates and their closest living relatives, animals. We were able to overcome initial failed efforts to establish genome editing in *S. rosetta* by engineering cycloheximide resistance in *rpl36a* as a selectable marker, similar to the use of selectable markers during the establishment of genome editing in other eukaryotes, including Fungi (Foster et al., 2018), green algae (Ferenczi et al., 2017), and nematodes (Arribere et al., 2014; Kim et al., 2014; Ward, 2015). Single-copy ribosomal protein genes like *rpl36a* offer certain advantages for engineering drug resistance markers with genome editing. First, resistance mutations in ribosomal protein genes have been genetically and biochemically characterized for a variety of drugs in diverse eukaryotes (Sutton et al., 1978; Ares and Bruns, 1978; Kawai et al., 1992; Dehoux et al., 1993; Kondo et al., 1995; Kim et al., 1998; Stevens et al., 2001; Garreau de Loubresse et al., 2014 and references therein). In our case, interpreting alignments among Rpl36a sequences from *S. rosetta* and organisms in the context of structures of eukaryotic ribosomes provided a starting point for customizing cycloheximide resistant alleles, a strategy that can also extend to other organisms. Second, the specificity of antibiotics that inhibit eukaryotic or prokaryotic translation can be leveraged to tailor genetic tools for particular organisms in complex communities. For example, cycloheximide binds selectively to eukaryotic ribosomes, resulting in the inhibition of *S. rosetta* growth and not that of its food source: live prey bacteria. Combining these advantages to establish genome editing in *S. rosetta* provided the first proof-of-principle for genome editing and allowed us to characterize the essential parameters before targeting other genes.

With the newfound potential for reverse genetics, we revisited the genetic basis of multicellular rosette development in *S. rosetta*. A previous forward genetic screen followed by mapping crosses implicated the C-type lectin gene *rosetteless* in the regulation of rosette development (Levin et al., 2014). At the time, however, it was not possible to independently corroborate *rosetteless* function with targeted mutations. In this study, we used genome editing to introduce a premature termination sequence in *rosetteless* and found that strains with the engineered *rosetteless* mutation have the same rosette defect phenotype as cells with the original *rtls^tl1^* mutation, demonstrating that *rosetteless* is necessary for rosette development. Moving forward, the approach established here will accelerate future forward genetic screens. It will now be possible for choanoflagellate researchers to introduce candidate mutations into a wild-type strain or correct the causative mutations in the original mutant strain to cleanly test the connection between genotype and phenotype.

Importantly, the establishment of genome editing in *S. rosetta* offers the first model choanoflagellate to investigate the ancestral and core functions of genes that evolved as integral regulators of animal biology. The *S. rosetta* genome (Fairclough et al., 2013) encodes receptors for immunity (e.g. Toll-like receptors), intercellular communication (e.g. receptor tyrosine kinases), and adhesion (e.g. cadherins, C-type lectins, and immunoglobulins) as well as master regulators of cell differentiation (i.e. forkhead, homeodomain, p53 and sox transcription factors). As a simple microbial model, *S. rosetta* now may serve as an accessible system for uncovering the conserved functions of genes that are not as readily studied in the more complex context of multicellular animals. Moreover, *S. rosetta* is just one tip on the choanoflagellate branch. A recent survey of 21 choanoflagellate transcriptomes revealed that choanoflagellates are at least as genetically diverse as animals (Richter et al., 2018), with other species retaining genetic pathways or exhibiting behaviors that are not found in *S. rosetta* (e.g., Marron et al., 2013; Leadbeater, 2015; Brunet et al., 2019). Together with future findings from *S. rosetta*, we anticipate that the establishment of genome editing in other choanoflagellates will provide an increasingly complete portrait of the last common ancestor of choanoflagellates and animals.

## Materials and Methods

### Culturing Choanoflagellates

Strains of *S. rosetta* were co-cultured with *Echinicola pacifica* bacteria (Levin and King, 2013; American Type Culture Collection [ATCC], Manassas, VA; Cat. No. PRA-390) in seawater-based media enriched with glycerol, yeast extract, and peptone to promote the growth of *E. pacifica* that serve as the choanoflagellate prey (Levin and King, 2013; Booth et al., 2018). We further supplemented this media with cereal grass (King et al., 2009; Fairclough et al., 2010; Carolina Biological Supply Company, Burlington, NC; Cat. No. 132375), which we call high nutrient media (Table S1), as we noticed that this addition promoted *S. rosetta* growth to a higher cell density (∼10^7^ cells/ml [Figure S4A] versus ∼10^6^ cells/ml [Booth et al., 2018]). To maintain rapidly proliferating cells in an abundance of nutrients, cultures were diluted 1 in 30 daily or 1 in 60 every two days into 6 ml of high nutrient media in 25 cm^2^ vented culture flasks (Corning, Oneonta, NY, USA; Cat. No. 430639) and incubated at 22°C and 60% relative humidity. To prevent an overgrowth of bacteria when *S. rosetta* experienced stress, such as after transfections or during clonal isolation, we cultured *S. rosetta* in low nutrient media, which is 0.375x high nutrient media (Table S1).

### Purification of outer membrane vesicles that contain RIFs

Rosette inducing factors (RIFs) contained in outer membrane vesicles (OMVs) from *Algoriphagus machipongonensis* (Alegado et al., 2013; ATCC; Cat. No. BAA-2233) can be used to induce rosette development in *S. rosetta* (Alegado et al., 2012; Woznica et al., 2016)*. A machipongonensis* OMVs were purified using the protocol in (Woznica et al., 2016). In summary, a 200 ml culture of 25x high nutrient media without cereal grass was inoculated from a single colony of *A. machipongonensis* and grown in a 1 l, baffled flask by shaking at 200 rpm for 3 days at 30°C. Afterwards, the bacteria were pelleted in 50 ml conical tubes by centrifugation at 4500*g* and 4°C for 30 min. The pellet was discarded and the supernatant was filtered through a 0.22 *µ*m vacuum filter. Outer membrane vesicles were pelleted from the filtered supernatant by ultracentrifugation at 36,000*g* and 4°C in a fixed-angle, Type 45 Ti rotor (Beckman Coulter Life Sciences, Indianapolis, IN; Cat. No. 339160) for 3 h. After discarding the supernatant, the pellet of outer membrane vesicles, which has an orange hue, was resuspended in a minimal volume of 50 mM HEPES-KOH, pH 7.5 and then incubated at 4°C overnight to fully dissolve the pellet. Last, the pellet was sterile filtered through a 0.45 *µ*m polyvinylidene fluoride syringe filter (Thermo Fisher Scientific, Waltham, MA; Cat. No. 09-720-4) into a sterile tube.

The rosette-inducing activity of the OMVs was tested by serially diluting the purified OMVs in a 24-well plate, with each well containing 0.5 ml of *S. rosetta* at a concentration of 10^4^ cells/ml and *E. pacifica*. The cells were incubated with OMVs at 22°C for 48 hours and then fixed with formaldehyde before counting the fraction of cells (*n* = 100) in rosettes. The dilution of lipids in which half of *S. rosetta* cells formed rosettes was defined as 2 unit/ml. All subsequent rosette inductions were performed with OMVs at a final concentration of 10 units/ml.

### Genome editing

Below we describe the considerations for the design and preparation of gRNAs and repair oligonucleotides for genome editing. The particular gRNAs and DNA repair template sequences for each given experiment are provided in Table S2.

#### Design and preparation of gRNAs

Upon inspecting the structure of the *Sp*Cas9 RNP poised to cleave a DNA target (Jiang et al., 2016), we concluded that sequences adjacent to and upstream of the PAM sequence (5’-NGG-3’), which have been reported to bias *Sp*Cas9 activity *in vivo* (Doench et al., 2014; Wu et al., 2014; Xu et al., 2015; Moreno-Mateos et al., 2015; Horlbeck et al., 2016; Liu et al., 2016; Kaur et al., 2016; Gandhi et al., 2017), likely influence *Sp*Cas9 recognition by stabilizing the conformation of the DNA target for cleavage. Therefore, we accounted for biases in *Sp*Cas9 recognition by choosing gRNAs sequences that conformed, as much as possible, to the motif 5’-HNNGR<u>SGG</u>H-3’, in which the PAM is underlined, N stands for any base, R stands for purine, S stands for G or C, and H stands for any base except G. This motif was first used to search for suitable targets (Peng and Tarleton, 2015) in cDNA sequences. We reasoned that initially searching for putative targets in cDNA sequences would ensure that gRNAs direct *Sp*Cas9 to cleave in protein coding regions of genes, and we later verified that putative gRNAs recognized genomic sequences instead of exon-exon junctions. Finally, we filtered out putative gRNA sequences with potential secondary sequences that can impede gRNA hybridization with DNA targets (Thyme et al., 2016) by evaluating their predicted secondary structures (Lorenz et al., 2011) and keeping gRNAs with predicted folding free energies greater than −1.5 kcal/mol.

gRNAs were prepared by annealing synthetic CRISPR RNA (crRNA) with a synthetic trans-activating CRISPR RNA (tracrRNA). The synthetic crRNA contains the 20 nt sequence for gene targeting and an additional sequence to anneal to the tracrRNA that binds to *Sp*Cas9. Alternatively, we also performed genome editing (Fig. 2 and Fig. S2H) with *in vitro* transcribed gRNAs (see supplementary information) that link the crRNA and tracrRNA into one continuous strand (Jinek et al., 2012; Chen et al., 2013), but we found that genome editing with crRNA/tracrRNA was the most time- and cost-effective. To prepare a functional gRNA complex from synthetic RNAs, crRNA and tracrRNA (Integrated DNA Technologies [IDT], Coralville, IA, USA) were resuspended to a final concentration of 200 *µ*M in duplex buffer (30 mM HEPES-KOH, pH 7.5; 100 mM potassium acetate; IDT, Cat. No. 11-01-03-01). Equal volumes of crRNA and tracrRNA stocks were mixed together, incubated at 95°C for 5 min in an aluminum heat block, and then the entire heat block was placed at room temperature to slowly cool the RNA to 25°C. The RNA was stored at −20°C.

#### Design and preparation of repair oligonucleotides

Repair oligonucleotides for generating knockouts were designed by copying the sequence 50 bases upstream and downstream of the *Sp*Cas9 cleavage site, which itself is 3 bp upstream of the PAM sequence (for example, 5’-N-cleave-NNN<u>NGG</u>-3’; PAM sequence underlined). A PTS (5’-TTTATTTAATTAAATAAA-3’) was inserted at the cleavage site. Importantly, this sequence has a stop codon (TAA) in each possible reading frame to terminate translate, a polyadenylation sequence (AATAAA) to terminate transcription, and a PacI sequence (5’-TTAATTAA-3’) that can be used to genotype with restriction digests. Moreover, the knockout sequence is palindromic, so it can be inserted in the sense or antisense direction of a gene and still generate a knockout.

Dried oligonucleotides (IDT) were resuspended to a concentration of 250 *µ*M in a buffer of 10 mM HEPES-KOH, pH 7.5, incubated at 55°C for 1 hour, and mixed well by pipetting up and down. The oligonucleotides were stored at −20°C.

#### Delivery of gene editing cargoes with nucleofection

*Sp*Cas9 RNPs and DNA repair templates were delivered into *S. rosetta* using a modified method for nucleofection (Booth et al., 2018). Here we describe here the complete transfection procedure and provide a publicly-accessible protocol specific for genome editing (https://dx.doi.org/10.17504/protocols.io.89fhz3n):

##### Cell Culture

Two days prior to transfection, 120 ml of high nutrient media was inoculated with a culture of *S. rosetta/E. pacifica* to a final concentration of *S. rosetta* at 8000 cells/ml. The culture was grown in a 3-layer flask (Corning; Cat. No. 353143), which has a surface area of 525 cm^2^, at 22°C and 60% humidity.

##### Assembly of Cas9/gRNA RNP

Before starting transfections, the *Sp*Cas9 RNP was assembled. For one reaction, 2 *µ*l of 20 *µ*M *Sp*Cas9 (NEB, Cat. No. M0646M or purified as described in supplementary information) was placed in the bottom of a 0.25 ml PCR tube, and then 2 *µ*l of 100 *µ*M gRNA was slowly pipetted up and down with *Sp*Cas9 to gently mix the solutions. The mixed solution was incubated at room temperature for 1 hour, which is roughly the time to complete the preparation of *S. rosetta* for priming (see below).

##### Thaw DNA oligonucleotides

Before using oligonucleotides in nucleofections, the oligonucleotides (prepared as above) were incubated at 55°C for 1 hour during the assembly of the *Sp*Cas9 RNP to ensure that they were fully dissolved.

##### Cell washing

*S. rosetta* cells were first prepared for nucleofection by washing away feeder bacteria. The 120 ml culture started two days previously was homogenized by vigorous shaking and then split into 40 ml aliquots in 50 ml conical tubes. The aliquots were vigorously shaken before centrifuging the cells for 5 min at 2000*g* and 22°C in a swinging bucket rotor. All but 2 ml of the supernatant, which remains cloudy with *E. pacifica* bacteria, was gently pipetted off of the pellet with a serological pipette; a fine tip transfer pipette gently removed the remaining liquid near the pellet. The three cell pellets were resuspended in artificial seawater (ASW; see table S1) for a total volume of 50 ml, combined into one conical tube, and vigorously shaken to homogenize the cells. For a second time, the resuspended cells were centrifuged for 5 min at 2000*g* and 22°C. The supernatant was removed as before, the pellet was resuspended in 50 ml of artificial seawater, and the cells were homogenized by vigorous shaking. The cells were centrifuged for a third time for 5 min at 2200*g* and 22°C. After removing the supernatant as described above, the cell pellet was resuspended in 400 *µ*l of ASW. The concentration of cells was determined by diluting 2 *µ*l of cells into 196 *µ*l of ASW. The diluted cells were fixed with 2 *µ*l of 37.5% formaldehyde, vortexed, and then pipetted into a fixed chamber slide for counting with Luna-FL automated cell counter (Logos Biosystems, Anyang, KOR; Cat. No. L20001). After determining the cell concentration, the washed *S. rosetta* cells were diluted to a final concentration of 5×10^7^ cell/ml and split into 100 *µ*l aliquots.

##### Priming

To prime *S. rosetta* cells for nucleofection, we treated them with a cocktail that removes the extracellular matrix as follows. Aliquots of washed cells were pelleted at 800*g* and 22°C for 5 min. The supernatant was gently removed with gel-loading tips and each pellet was resuspended in 100 *µ*l of priming buffer (40 mM HEPES-KOH, pH 7.5; 34 mM lithium citrate; 50 mM l-cysteine; 15% [wt/vol] PEG 8000; and 1 μM papain [Millipore Sigma, St. Louis, MO; Cat. No. P3125-100MG]). After incubating cells for 30-40 min, 10 *µ*l of 50 mg/ml bovine serum albumin was added to each aliquot of primed cells to quench proteolysis from the priming buffer. Finally, the cells were centrifuged at 1250g and 22°C for 5 min, the supernatant was removed, and the pellet was resuspended in 25 *µ*l of SF Buffer (Lonza, Basel, Switzerland; Cat. No. V4SC-2960). The resuspended cells were stored on ice while preparing nucleofection reagents.

##### Nucleofection

Each nucleofection reaction was prepared by adding 16 *µ*l of ice-cold SF Buffer to 4 *µ*l of the *Sp*Cas9 RNP that was assembled as described above. (For reactions that used two different gRNAs, each gRNA was assembled with *Sp*Cas9 separately and 4 *µ*l of each RNP solution was added to SF buffer at this step). 2 *µ*l of the repair oligonucleotide template was added to the *Sp*Cas9 RNP diluted in SF buffer. Finally, 2 *µ*l of primed cells were added to the solution with *Sp*Cas9 RNP and the repair template. The whole solution, which has a total volume of 24 *µ*l (30*µ*l for two different *Sp*Cas9 RNPs and repair templates), was placed in one well of a 96-well nucleofection plate. The well was pulsed in a Lonza shuttle nucleofector (Lonza, Cat. No. AAF-1002B and AAM-1001S) with the CM156 pulse.

##### Recovery

Immediately after transfection, 100 *µ*l of ice-cold recovery buffer (10 mM HEPES-KOH, pH 7.5; 0.9 M sorbitol; 8% [wt/vol] PEG 8000) was added to each transfection and gently mixed by firmly tapping the side of the plate or cuvette. After the cells rested in recovery buffer at room-temperature for 5 min, the whole volume of a nucleofection well was transferred to 2 ml of low nutrient media in one well of a 6 well plate. After 30 min, 10 *µ*l of 10 mg/ml *E. pacifica* (prepared by resuspending a frozen 10 mg pellet of *E. pacifica* in ASW) was added to each well and the 6 well plate was incubated at 22°C and 60% relative humidity for downstream experiments.

### Establishing Clonal Strains

Here we describe how to isolate clones to establish strains. For a complete list of strains used in this study, see Table S3.

#### Cycloheximide Selection

One day after transfecting *S. rosetta* with *Sp*Cas9 RNPs repair oligonucleotides for *rpl36a^P56Q^* (Fig. 2), 10 *µ*l of 1 *µ*g/ml cycloheximide was added to a 2 ml culture of transfected cells. The cells were incubated with cycloheximide for 5 days prior to genotyping and clonal isolation.

#### Clonal Isolation

To prepare cells for clonal isolation by limiting dilution, the initial cell density was determined by fixing a 200 *µ*l sample of cells with 5 *µ*l of 37.5% (w/v) formaldehyde and then by counting the fixed cells with a hemocytometer (Hausser Scientific, Horsham, PA; Cat. No. 1475) or Luna-FL automated cell counter. The cells were by diluted to a final concentration of 3 cells/ml in low nutrient sea water and then distributed in a 96 well plate with 100 *µ*l/well. Thus, the mean frequency of finding a cell in each well is 0.3, which, according to a Poisson distribution, corresponds to a >99% probability that a given well with *S. rosetta* was founded from a single cell. Cells were grown in a 96 well plate for 5-7 days at 22 °C and 60% relative humidity. Lastly, the plate was screened using phase contrast microscopy to identify wells with *S. rosetta*. Finally, larger cultures of high nutrient media were inoculated with clonal isolates to establish strains.

### Genotyping by Sanger Sequencing (Fig. 2-3 and S2-S3)

Cells were harvested for genotyping by centrifuging 1 ml of cells at 4250*g* and 22°C for 5 min. The supernatant was removed with a fine tip transfer pipette. (Optional: To remove lingering DNA from cells that die in the course of cycloheximide selection, the pellet was resuspended in 50 *µ*l DNase buffer [10 mM Tris-HCl, pH 7.6; 1 M sorbitol; 2.5 mM magnesium chloride; 0.5 mM calcium chloride; 0.1 U/*µ*l Turbo DNase (Thermo Fisher Scientific; Cat. No. AM2238)] and incubated at room temperature for 30 min. Afterwards, the cells were centrifuged as before, discarding the supernatant.) The cell pellet was dissolved in 100 *µ*l of DNAzol Direct (20 mM potassium hydroxide, 60% [w/v] PEG 200, pH 13.3-13.5; Molecular Research Center, Inc., Cincinnati, OH; Cat. No. DN131). 5 *µ*l of the dissolved cells were added to a 50 *µ*l PCR reaction (Q5 DNA polymerase, NEB; see Table S2 for primer sequences) and amplified with 36 rounds of thermal cycling. Samples dissolved in DNAzol direct can be directly added to PCR reactions because the pH of DNAzol Direct dramatically drops upon a ten-fold or greater dilution [Chomczynski and Rymaszewski, 2006]. The PCR product was purified using magnetic beads (Oberacker et al., 2019) and then submitted for Sanger sequencing (UC Berkeley DNA Sequencing facility).

### Cell proliferation assays (Fig. S4)

We characterized the cell proliferation rates of *S. rosetta* strains by monitoring the concentration of cells over time to fit logistic growth curves and determine the doubling time. Cell proliferation assays started by diluting cultures to a concentration of 10^4^ cells/ml in high nutrient media and then distributing 0.5 ml of culture into each well of a 24 well plate. Every ∼12 h, the entire contents of one well were thoroughly homogenized by pipetting up and down and then transferred to a 1.5 ml conical tube. Three independent wells were taken for triplicate measures of cell concentration at every time point. The cells were fixed with 20 *µ*l of 37.5% formaldehyde and mixed by vortexing. The fixed cells were stored at 4°C until the sample was used for determining the cell concentration after the full growth course.

The cell concentration was determined by counting the number of cells in a fixed-volume imaging chamber. In detail, the fixed cells were thoroughly homogenized by vortexing for 10 s and then pipetted up and down before transfer into a chamber of a Smart Slide (ibidi USA, Inc., Firchburg, WI; Cat. No. 80816) that has a fixed height of 200 *µ*m. After allowing cells to settle to the bottom for 5 min, each chamber was imaged on an Axio Observer.Z1/7 Widefield microscope (Carl Zeiss AG, Oberkochen, Germany) and recorded with a Hamamatsu Orca-Flash 4.0 LT CMOS Digital Camera (Hamamatsu Photonics, Hamamatsu City, Japan) using either phase contrast for 10x (objective), in-focus images or a 20x brightfield image with a 1 *µ*m overfocus to make the cells appear dark on a light gray background. The volume for each image was calculated from the image area, which was calibrated on the microscope, and the fixed height of the imaging chamber: 3.54×10^−4^ ml when imaged at 10x and 8.86×10^−5^ ml when imaged at 20x. Using automated particle detection in Fiji (Schindelin et al., 2012), cells were counted in each 20x image by thresholding the image to make cells appear as black spots on a white background and then each circular spot was counted with the “Analyze Particles” function. For early time points with fewer numbers of cells, we manually counted cells in 10x images to include more cells in a greater area for a more accurate count.

Each time course was fit by least absolute deviation curve fitting to the logistic equation:

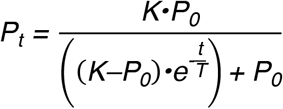

where *P_t_* is the cell density at time (*t*), *K* is the carrying capacity, *P_0_* is the initial cell density, and *T* is the doubling time.

### Live-cell Microscopy (Figs. 1 and 3)

Glass-bottomed dishes (World Precision Instruments, Sarasota, FL; Cat. No. FD35-100) were prepared for imaging by covering the bottom with 500 *µ*l of 0.1 mg/ml poly-D-lysine (Millipore Sigma; Cat. No. P6407-5MG) and incubating for 15 min. The poly-D-lysine was removed and then the dish was washed three times with 500 *µ*l of ASW. Cells were placed into the dish by gently pipetting 500 *µ*l of cells with a wide pipette tip.

Differential interference contrast (DIC) microscopy images were captured with a Zeiss Axio Observer.Z1/7 Widefield microscope with a Hamamatsu Orca-Flash 4.0 LT CMOS Digital Camera (Hamamatsu Photonics, Hamamatsu City, Japan) and 40×/NA 1.1 LD C-Apochromatic water immersion, 63×/NA1.40 Plan-Apochromatic oil immersion, or 100× NA 1.40 Plan-Apochromatic oil immersion objectives (Zeiss).

### Immunofluorescent Staining and Imaging (Figs. 1 and S3)

200 *µ*l of *S. rosetta* cells were gently pipetted into chamber slides (ibidi; Cat. No.80826) coated with poly-D-lysine (see live cell imaging for coating procedure). Importantly, cells were pipetted using a tip that had been trimmed to create a larger bore for reducing shear forces. The cells were incubated on the coverslip for 30 min to allow the cells to adsorb to the surface.

Cells were fixed by adding 200 *µ*l of 6% acetone in cytoskeleton buffer (10 mM MES, pH 6.1; 138 KCl, 3 mM MgCl_2_; 2 mM ethylene glycol-bis(β-aminoethylether)-*N*,*N*,*N*’,*N*’-tetraacetic acid [EGTA]; 600 mM sucrose) and then incubated for 10 min at room temperature. After removing 200 *µ*l from the chamber, 200 *µ*l of 4% formaldehyde diluted in cytoskeleton buffer was added to the chamber and then incubated for 15 min at room temperature. Last, the coverslip was gently washed three times with 200 *µ*l of cytoskeleton buffer.

Cells were permeabilized by washing the coverslip once with 200 *µ*l of permeabilization buffer (PEM [100 mM PIPES, pH 6.95; 2 mM EGTA; 1 mM MgCl_2_] with 1% [wt/vol] bovine serum albumin (BSA)-fraction V and 0.3% [vol/vol] Triton X-100) and then incubated for 60 min upon a second addition of permeabilization buffer. Afterwards, 200 *µ*l of the permeabilization buffer was replaced with primary antibodies diluted in permeabilization buffer, 1 *µ*g/ml mouse DM1A anti-*α*-tubulin antibody (Thermo Fisher Scientific; Cat. No. 62204) and 1:200 rabbit anti-Rosetteless (Levin et al., 2014). After the samples were incubated in primary antibody for 2 h, the chamber was gently washed three times with 200 *µ*l permeabilization buffer. Next, 200 *µ*l of permeabilization buffer with 10 *µ*g/ml donkey anti-mouse immunoglobulin G–AlexaFluor568 (Thermo Fisher Scientific; Cat. No. A10037), donkey anti-rabbit immunoglobulin G–AlexaFluor647 (Thermo Fisher Scientific; Cat. No. A32795), 10 μg/ml Hoechst 33342 (Thermo Fisher Scientific; Cat. No. H3570), and 4 U/ml Phalloidin-AlexaFluor488 (Thermo Fisher Scientific; Cat. No. A12379) was added to the chamber and then incubated for 40 min. Afterwards, the chamber was washed five times with PEM.

Immunostained samples were imaged on a Zeiss Axio Observer LSM 880 with an Fast Airyscan detector and a 40x/NA1.1 Plan-Apochromatic water immersion objective (Zeiss) by frame scanning in the superresolution mode with the following settings: 50 × 50 nm pixel size; 220 nm *z*-step; 0.73 μs/pixel dwell time; 750 gain; 488/561/633 nm multiple beam splitter; 633-nm laser operating at 16% power with a 570-620/645 bandpass/longpass filter; 561-nm laser operating at 16% power with a 570-620/645 bandpass/longpass filter; 488-nm laser operating at 14% power with a 420-580/495-550 bandpass filters.

### Next-generation sequencing (Fig. 4 and Fig. S5)

We performed deep sequencing of edited cells to quantify the efficiency of genome editing (Fig. 3 and Fig. S5). The transfections were performed as above with the following modifications: Two transfections were conducted for each condition and combined into 1 ml of low nutrient media (see Table S1 for recipe). One day after transfection, the cells were harvested and dissolved in 50 *µ*l of DNAzol direct (Molecular Research Center, Inc.). Three independent transfections performed on different days provided replicate measures for each condition (Fig. 3C and Fig. S5B).

To preserve the diversity of sequences during PCR, six parallel PCR reactions (Q5 DNA polymerase, NEB) were set up with 30 *µ*l of sample. The target locus was amplified in 15 thermal cycles, purified using magnetic beads (UC Berkeley DNA sequencing facility), and pooled together in a total volume of 180 *µ*l. Importantly, the primers for this first round of PCR had a randomized 6-nucleotide sequence in the forward primer to distinguish PCR duplicates (primer sequences in Table S2), which allowed us to identify 4096 unique sequences. Extending this randomized sequence would result in higher sensitivity.

A second round of PCR was performed to attach adapters for Illumina sequencing. For these reactions, four replicate PCR reactions were set up with 25 *µ*l of the purified products from the first round of PCR and primers with sequencing adapters and unique sample barcodes were attached in 5 thermal cycles. Afterward, the PCR products were purified using magnetic beads (UC Berkeley DNA sequencing facility) and their quality was assessed on a Bioanalyzer (UC Berkeley Functional Genomic Laboratory). The bioanalyzer traces showed that the amplicons were the proper size, yet a similar concentration of residual PCR primers remained in each sample. After quantifying DNA (Qubit; Thermo Fisher Scientific) and pooling equimolar amounts of sample, the amplicons were further purified with magnetic beads (UC Berkeley Functional Genomics Lab) and the concentration was verified using qPCR. The library was sequenced on a miSeq sequencer (Illumina, San Diego, CA) using the V3 chemistry (Illumina) for 300 rounds of paired-end sequencing, which gives up to 600 bases of sequence per sample. After sequencing the samples were separated based on their unique barcodes for further analysis of individual samples.

The editing efficiency for each sample was calculated from high-quality, unique reads. First, we used tools from the Galaxy Project (Afgan et al., 2018) to join paired-end reads into one read (fastq-join) and then retain high quality sequences (Galaxy–Filter by quality: 100% of bases with quality scores ≥ 30) with 50 bp of expected sequence from the *rosetteless* locus on the 5’ and 3’ ends of the amplicon (Galaxy–Cutadapt: 50 base overlap with 0.1 maximum error rate). Next, the reads were filtered for unique instances of the randomized barcode sequence from the first round PCR primers (Galaxy–Unique). We then combined matching amplicon sequences into unique bins, while counting the number of sequences in each bin (Galaxy–Collapse). The FASTA file of aligned sequences (Galaxy–ClustalW) from this initial processing was further analyzed using a custom script (Supplementary File S1). To quantify the instances of template-mediated repair, we counted the number of sequences that had the PTS. Untemplated mutations were counted from insertions and deletions larger than 1 bp. The remaining sequences, those that were the same length as the *rosetteless* locus but did not have the exact amplicon sequence, were counted as single-nucleotide polymorphisms (SNPs). The outputs from each category were also visually inspected to reclassify incorrect calls, such as a few instances of template-directed repair in the insertions and deletion category due to mutations in the PTS. The SNP data revealed that conditions with or without the addition of *Sp*Cas9 or repair templates had same SNP frequency. Therefore, we only compared reads categorized as template-mediated repair or untemplated insertions and deletions. Importantly, this analysis may overlook some instances where DNA repair resulted in sequences that maintained the original sequence or introduced SNPs, thereby underestimating the efficiency of non-templated repair.

## Acknowledgements

We thank the following people for insights and support that helped advance this work: Heather Szmidt-Middleton, Laura Wetzel, Monika Sigg, Lily Helfrich, Arielle Woznica, Sabrina Sun, Tara DeBoer, Jorge Santiago-Ortiz, and Kayley Hake helped with and provided feedback on early experiments. Through the Gordon and Betty Moore Foundation Marine Microbiology Initiative (GBMF MMI), Manny Ares first brought cycloheximide resistance alleles to our attention. The following people stimulated helpful discussions: Fyodor Urnov, Stephen Floor, Chris Richardson, Jacob Corn, David Schaffer, Niren Murthy, Craig Miller, and members of the King Lab. Brett Stahl, Shana McDevitt, the UC Berkeley Vincent Coates Sequencing Center, and the UC Berkeley Functional Genomics Laboratory provided help with sequencing. This work was supported, in part, by a GBMF MMI Experimental Model Systems grant. DSB was supported through a Simons Foundation Fellowship from the Jane Coffin Childs Memorial Fund for Medical Research.

## Author Contributions

DSB and NK conceived of the project and wrote the manuscript. DSB designed, performed and interpreted experiments.

## Supplementary Information

### Supplementary Materials and Methods

#### *In vitro* transcription of gRNAs

DNA templates for *in vitro* transcription of gRNAs were amplified by PCR (Q5 DNA Polymerase; New England Biolabs [NEB], Ipswich, MA, USA, Cat. No. M0491L) from synthetic DNA templates (IDT; Table S2) that had a T7 promoter sequence appended to the 5’ end of the guide sequence and a trans-activating CRISPR RNA (tracrRNA) sequence (Chen et al., 2013) at the 3’ end. The purified DNA templates (PCR cleanup kit; Qiagen, Venlo, NLD; Cat. No. 28006) were used to synthesize gRNAs with T7 RNA polymerase (Milligan and Uhlenbeck, 1989) in reactions set up with these components: 40 mM Tris-HCl, pH 8.0; 2.5 mM spermidine; 0.01% (v/v) Triton X-100; 5 mM GTP; 5 mM UTP; 5 mM ATP; 5 mM CTP; 80 mg/ml PEG 8000; 32 mM magnesium chloride; 5 mM dithiothreitol; 10 ng/*µ*l template DNA; 0.5 U/*µ*l SUPERase•In (Thermo Fisher Scientific, Waltham, MA; Cat. No. AM2696); 2 U/*µ*l T7 RNA polymerase (Thermo Fisher Scientific, Cat. No. EP0113); 0.025 mg/ml pyrophosphatase (Thermo Fisher Scientific, Cat. No. EF0221). After incubating the transcription reaction at 37°C for >4 h, the DNA template was digested with the addition of 0.1 U/*µ*l TURBO DNase (Thermo Fisher Scientific, Cat. No. AM2239). After assessing the transcription products on denaturing, urea-polyacrylamide gel electrophoresis (PAGE), we found that the *in vitro* transcriptions yielded high amounts of gRNA with few byproducts. Therefore, we used a simplified protocol to purify gRNAs by first removing contaminating nucleotides with a desalting column (GE Healthcare Lifesciences, Pittsburgh, PA; Cat. No. 17085302) to exchange gRNA into 1 mM sodium citrate, pH 6.4. The gRNAs were then precipitated from the solution by adding 0.25 volumes of RNA precipitation buffer (1.2 M sodium acetate, pH 5.2; 4 mM EDTA-NaOH, pH 8.0; 0.04% sodium dodecyl sulfate [SDS]) and 2.5 volumes of ethanol. The precipitated RNA was centrifuged for 60 min at 4°C, washed once with 70% ethanol/water, and finally resuspended in 1 mM sodium citrate, pH 6.4.

After determining the concentration of gRNA, which has a 260 nm extinction coefficient of 1.41×10^6^ M^−1^cm^−1^, by UV-vis spectroscopy, the gRNA was diluted to a final concentration of 50 *µ*M with 1 mM sodium citrate, pH 6.4. To ensure that the gRNA was properly folded, the gRNA was placed at 95°C for 5 min in an aluminum heat block and then slowly cooled to 25°C by placing the aluminum block on a room temperature bench top. Finally, gRNAs were stored at −20°C.

#### *Sp*Cas9 expression and purification

For efficient genome editing, we purified or purchased (NEB, Cat. No. M0646M) an engineered version of *Streptomyces pyogenes* Cas9 that has SV40 nuclear localization sequences (NLS) at the amino- and carobxy-termini of *Sp*Cas9. Below we describe a simplified purification procedure based on the previously published work (Jinek et al., 2012).

##### Vector construction

Using a variation of Gibson cloning (Gibson et al., 2009; NEB, Cat. No. E2621L), we modified a vector (Jinek et al., 2012; Addgene, Watertown, MA; Cat. No. 69090) for expressing *Sp*Cas9 in *Escherichia coli* by inserting tandem SV40 NLSs at the amino terminus of *Sp*Cas9. A similar construct (Addgene, Cat. No. 88916) has been shown to increase nuclear localization in mammalian cells (Cong et al., 2013; Staahl et al., 2017). The expression vector has a hexahistidine (His_6_) tag and maltose binding protein (MBP) fused to the amino terminus of *Sp*Cas9. A tobacco etch virus (TEV) protease cleavage site between MBP and the amino terminal nuclear localization sequence on Cas9 facilitates the removal of the His_6_-MBP tag from *Sp*Cas9.

##### Protein expression

The *Sp*Cas9 expression vector was transformed into the BL21 Star (DE3) strain of *E. coli* (Thermo Fisher Scientific, Cat. No. C601003), and a single colony of the transformants was inoculated into Miller’s LB broth (Atlas, 2010) for growing a starter culture overnight at 37°C with shaking at 200 rpm. 20 ml of the starter culture was diluted into 1 L of M9 medium (Atlas, 2010) and the culture was grown at 37°C with shaking at 250 rpm until the OD_600_ = 0.60. At that cell density, the culture was shifted to 16°C for 15 min and *Sp*Cas9 expression was induced by addition isopropyl β-D-1-thiogalactopyranoside (IPTG) to a final concentration of 0.5 mM. The culture was grown at 16°C overnight and cells were harvested by centrifugation at 4900*g* and 4° for 15 min in a swinging bucket centrifuge. The supernatant was discarded and the bacterial pellets were flash frozen in liquid nitrogen and stored at −80°C.

##### Protein Purification

The bacterial pellet was lysed by resuspending 1g of bacterial pellet in 9 ml of lysis buffer (150 mM potassium phosphate, pH 7.5; 500 mM sodium chloride; 5 mM imidazole, pH 8.0; 1 mM Pefabloc SC; 2 mM 2-mercaptoethanol; 10% [v/v] glycerol; 1 protease inhibitor tablet [cOmplete, EDTA-free; Roche; Cat. No, 04693132001] per 20 ml of lysate) and lysing with a microfluidizer. The lysate was centrifuged at 30,000*g* and 4°C for 30 min to remove insoluble debris.

The supernatant was passed through an Ni-NTA Agarose column (Qiagen, Cat. No. 30210) equilibrated in elution buffer (150 mM potassium phosphate, pH 7.5; 500 mM sodium chloride; 5 mM imidazole, pH 8.0; 2 mM 2-mercaptoethanol; 10% [v/v] glycerol), using 1 ml of resin per 10 grams of bacterial pellet. The column was washed with 10 column volumes (CV) of lysis buffer, 5 CV of elution buffer supplemented with 10 mM imidazole, and 3 CV of wash elution buffer supplemented with 20 mM imidazole. The protein was eluted from the column with 4 CV of elution buffer supplemented with 240 mM imidazole. After determining the protein concentration by UV-vis spectroscopy (using an extinction coefficient of 0.18829 *µ*M^−1^cm^−1^ for *Sp*Cas9), TEV protease was added at 1:20 molar ratio of TEV protease to *Sp*Cas9. SpCas9 supplemented with TEV protease was placed in a dialysis bag with a 3500 dalton molecular weight cut-off (MWCO) and dialyzed against dialysis buffer (100 mM potassium phosphate, pH 7.5; 2 mM 2-mercaptoethanol; 10 mM imidazole; 10% [v/v] glycerol) overnight at 4°C. Afterwards, the dialyzed protein was passed over the Ni-NTA column that had been equilibrated in dialysis buffer to remove His_6_-MBP tag from *Sp*Cas9, which is in the flow through. The flow through was loaded onto HiTrap SP High-Performance (GE Healthcare Lifesciences, Cat. No. 17-1152-01) column that had been equilibrated in dialysis buffer. The column was extensively washed with dialysis buffer prior to eluting the protein in S-elution buffer (500 mM potassium phosphate, pH 7.5; 3 mM dithiothreitol; 0.3 mM EDTA-KOH, pH 8.0; 10% [v/v] glycerol). The purity was evaluated by SDS-PAGE, and the concentration was measured using UV-vis spectroscopy. Afterwards, the purified protein was concentrated with a 100,000 MWCO centrifugal filter to a final concentration of 20-25 *µ*M. The concentrated protein was flash-frozen in liquid nitrogen and stored at −80°C.

**Figure S1:**
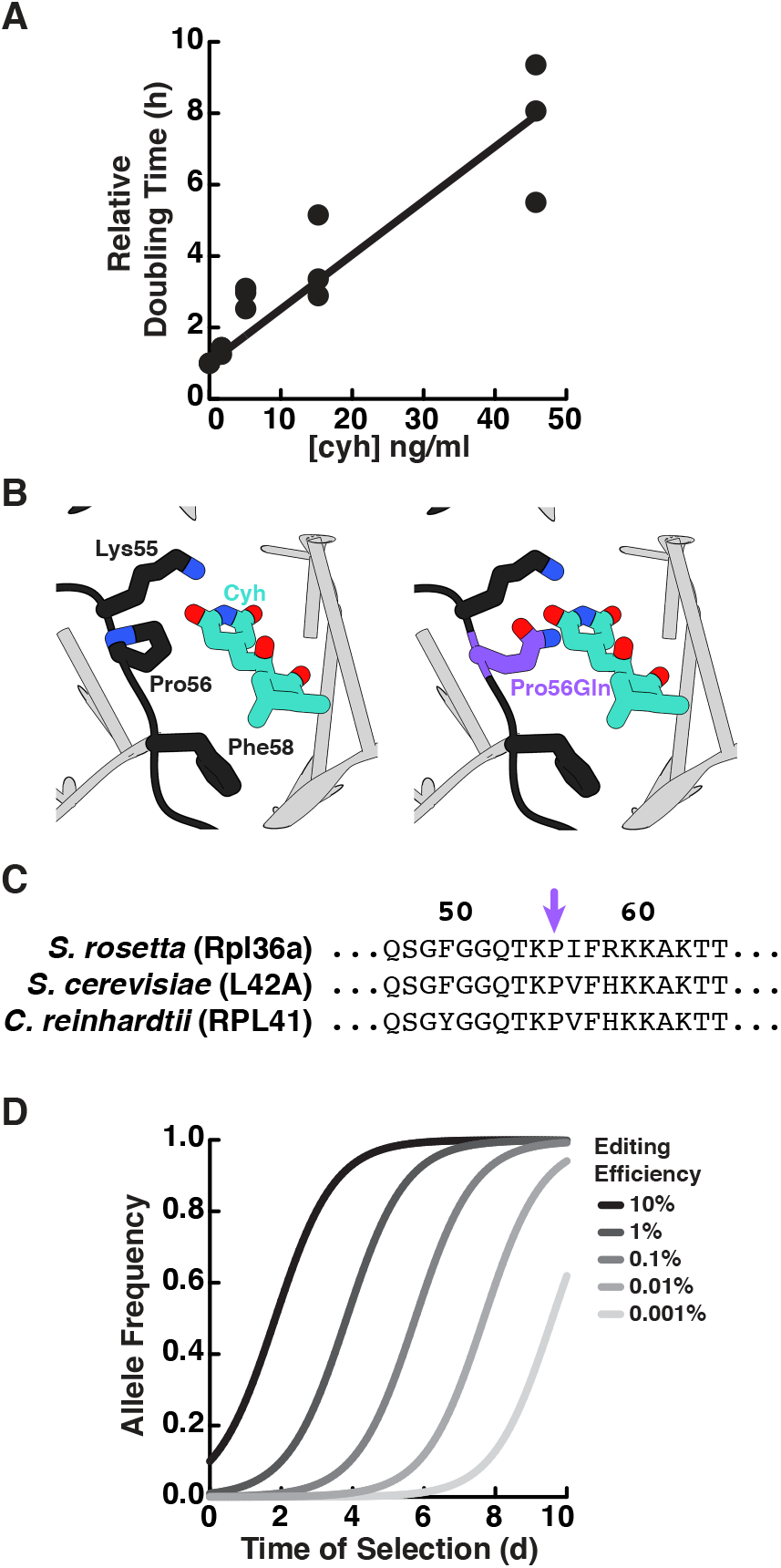
An approach for selecting cycloheximide resistance in *S. rosetta*. (A) Cycloheximide inhibits *S. rosetta* growth. We seeded each well of a 24-well plate with 0.5 ml of cells at 2×10^4^ cells/ml in a 3-fold serial dilution of cycloheximide, including a condition without cycloheximide. Three independent wells were set up for each concentration of cycloheximide for replicate measures. The cell density at each concentration was determined after 48 h by counting with a hemocytometer. To establish a relationship between growth rate and cycloheximide concentration, the cell density was transformed into relative growth rates (*S*) using a rearranged form of the logistic equation:

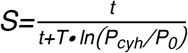

in which the cell density in a given cycloheximide concentration (*P_cyh_*) at time (*t* = 48 h) was normalized by the cell density without cyloheximide (*P_0_*) and the doubling time without cycloheximide (*T* = 10 h) was taken from growth curves of wild-type strains (Fig. S4A). After performing a linear fit of the data, we determined that 5 ng/ml of cycloheximide retards the growth rate two-fold. (B) The cycloheximide resistant mutation *rpl36a^P56Q^* disrupts cycloheximde binding to the large ribosomal subunit of yeast (left). A crystal structure of cycloheximide bound to the yeast 80S ribosome (Garreau de Loubresse et al., 2014; PDB 4U3U) shows that the yeast ortholog of Rpl36a (L42A; black) and ribosomal RNA (gray) form the cycloheximide binding pocket. The most critical residues in L42A for cycloheximide binding are Lys55, Pro56, and Phe58 (*right*). *In silico* modeling (Goddard et al., 2005) of cycloheximide resistance mutations (Bae et al., 2018) shows that some rotamers of the Pro56Gln substitution (purple) disrupt van der Waals interactions and cause steric clashes. (C) The *S. rosetta* ortholog of Rpl36a conserves residues that bind cycloheximide in yeast and *Chlamydomonas rheinhardtii*. A sequence alignment (Sievers et al., 2011) of Rpl36a orthologs from *S. rosetta*, *S. cerevisiae* (L42A), and *C. rheinhardtii* (RPL41) shows that *S. rosetta* conserves residues for cycloheximide binding, the most critical of which is Pro56 (purple arrow). (D) The efficiency of genome editing alters the selection of edited alleles. Using growth parameters determined from *S. rosetta* growth curves (Fig. S4) and cycloheximide inhibition (Fig. S1A), we modeled the selection for cycloheximide resistance using the following equation:

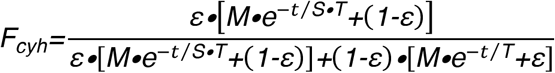

where *F_cyh_* is the frequency of the cycloheximide resistant allele, *ε* is the genome editing efficiency (which also corresponds to the initial frequency of the edited allele), *M* is the ratio of the carrying capacity to the cell density (which we set as an arbitrarily large number because continuous passaging in the laboratory can keep the cell population far from the carrying capacity), *t* is the time of growth after starting selection, *T* is the doubling time in the absence of selection (which is 10 h, see Fig. S4), and *S* is the relative growth rate in the presence of selection (which we set to 2, based on the relative growth rate upon adding 5 ng/ml of cycloheximide to cells (panel A). Notably, this model only captures the relative changes in growth upon selection and assumes that the edited allele is insensitive to the drug; the model does not include a term for the rate of cells dying, which we observed to happen after 3 days of selection. Nonetheless, this model helped us determine that after 5 days of growth in a selective media, we could expect to observe genome editing events occurring at frequencies >0.01%.

**Figure S2:**
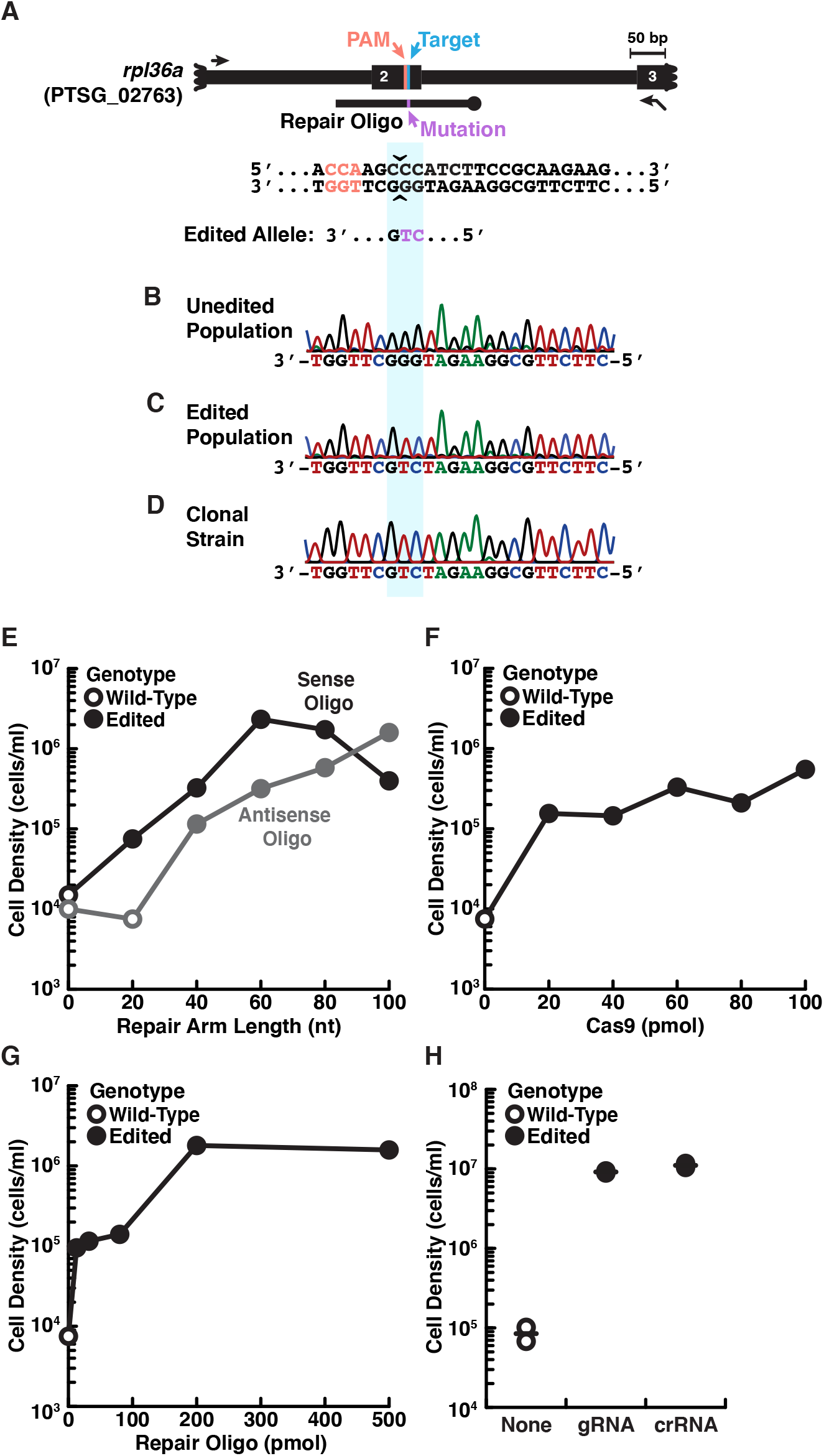
Engineered cycloheximide resistance establishes genome editing conditions. (A) The design of a cycloheximide resistant allele, *rpl36a^P56Q^*, in *S. rosetta*. The protospacer adjacent motif (PAM, orange) next to the 56^th^ codon of *rpl36a* (Target, cyan), which is located on the second exon (thick black line labeled 2), provides a suitable site to design a gRNA that targets *Sp*Cas9 cleavage (sequence is shown underneath the locus schematic, and carets indicate the target cleavage site). A repair oligonucleotide (black line with knob) introduces a cycloheximide resistant allele, *rpl36a^P56Q^* (Mutation, purple), flanked by 100 bases of homologous sequence. The sequence of the edited allele is shown below. (B-D) A comparison of genotypes from populations of unedited cells (B), edited cells (C), and a strain established from a clonal isolate of edited cells (D) shows that cycloheximide selection enriches for *rpl36a^P56Q^*. The genotype for each population was determined by amplifying the locus with primers surrounding the editing site (black arrows in panel A) that did not overlap in sequence with the repair oligonucleotide. One of the primers had a T3 primer binding site for Sanger sequencing of amplicons (black arrow with flap). Remarkably, after selection, the wild-type allele was not detected (B). (E) *S. rosetta* uses repair oligonucleotides with >20 nt homology arms for genome editing. Truncations of repair oligonucleotides encoding the *rpl36a^P56Q^* allele were designed in the same orientation as gRNAs (sense, black dots and lines) or the opposite orientation (antisense, gray dots and lines). 24 h after *S. rosetta* recovered from transfections with repair templates and *Sp*Cas9 RNPs, cycloheximide was added to grow cells in selective media for five days, at which time the cells were harvested for counting cell density and for genotyping. Closed circles indicate that the consensus genotype of the cell population had the *rpl36a^P56Q^* allele in Sanger sequencing; whereas, open circles indicate that the cell population had the wild-type allele. The results are from one of two independent experiments. Notably, we observed a slight bias for repair oligonucleotides in the sense direction, particularly with shorter homology arms of 20 bases. Because repair templates in the sense orientation with 40-80 bases of homologous sequence resulted in the best editing, we performed subsequent optimization with a sense repair oligonucleotide that 50-base homology arms on each side of the double-stranded break. (F) Small quantities of *Sp*Cas9 RNPs are sufficient to initiate genome editing. Decreasing concentrations of *Sp*Cas9 RNP (*Sp*Cas9 was the limiting factor) and a constant amount of repair template were transfected into *S. rosetta*. After characterizing genome editing outcomes by counting cell density and sequencing the consensus genotype (described in panel E), we found that low concentrations of *Sp*Cas9 (20 pmol) were sufficient to introduce the *rpl36a^P56Q^* allele. Results are from one of two independent experiments. (G) High concentrations of repair oligonucleotides increase genome editing efficiency. A serial dilution of a repair template was delivered into *S. rosetta*. The cell density and consensus genotypes from these experiments show that all concentrations of repair template can introduce the *rpl36a^P56Q^* allele, but the higher cell densities recovered after transfection with increasing concentrations of repair templates indicate more efficient editing. The data are from one of two independent experiments. (H) The addition of gRNAs stimulates genome editing. Genome editing was performed by delivering a repair oligonucleotide with *Sp*Cas9 without the addition of any gRNA or with a gRNA that was prepared from in vitro transcriptions (noted as gRNA in figure) or with a synthetic crRNA that was annealed to a synthetic tracrRNA (noted as crRNA). The consensus genotype and cell densities from these experiments show that gRNAs are necessary for editing and that gRNAs from either source were sufficient for editing. The dots show two independent experiments and lines show their average result.

**Figure S3:**
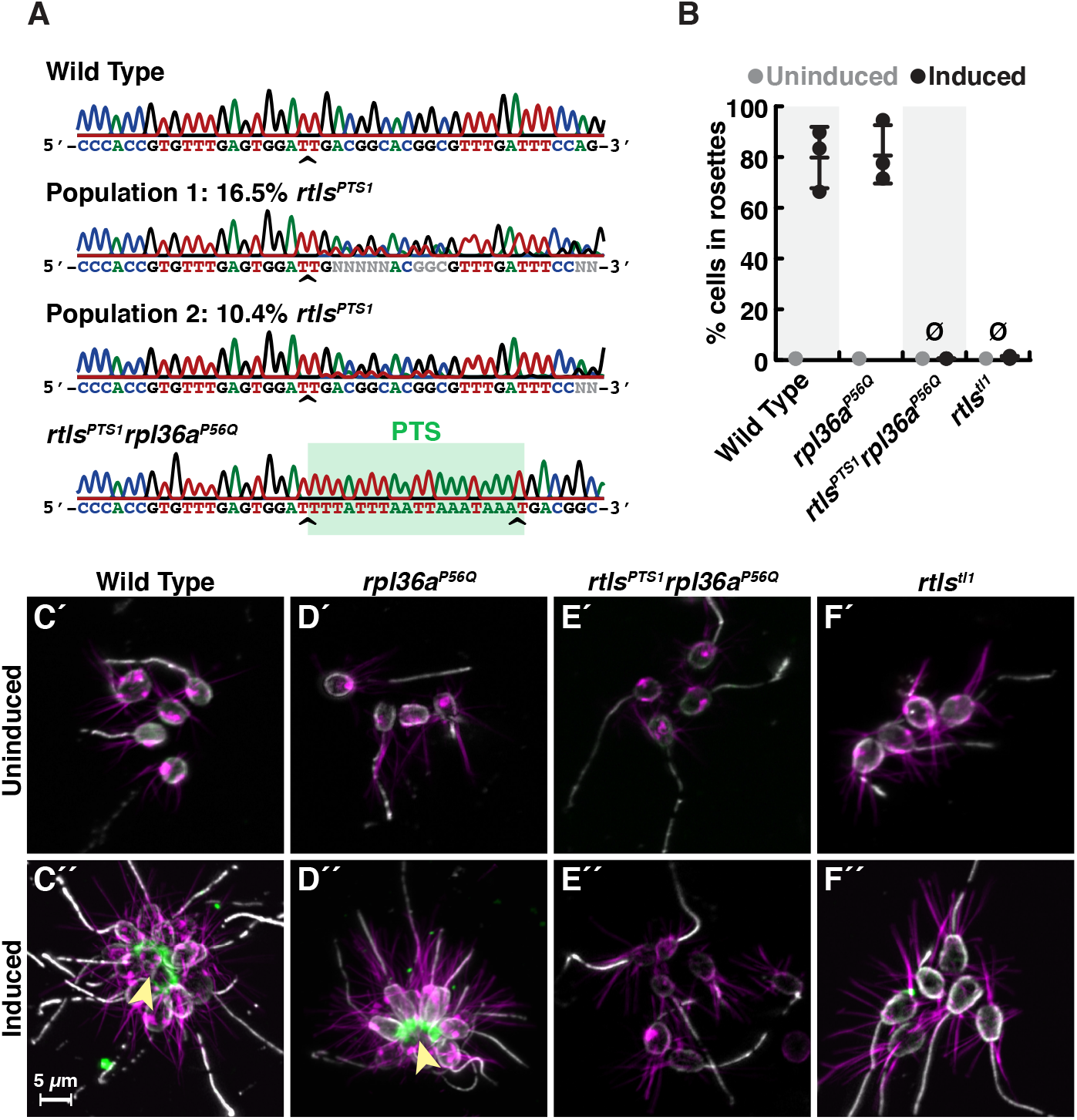
Phenotypes of *rosetteless* mutants correspond to their genotypes. (A) The consensus genotype at the site of *rosetteless* editing in cell populations selected for cycloheximide resistance indicates the presence of the *rtls^PTS1^* allele. In a wild-type strain (top) and a clonal isolate of *rlts^PTS1^ rpl36a^P56Q^* (bottom), Sanger sequencing at the *rtls* exon 4 reveals no heterogeneity in sequence. In populations of cells that had been co-edited to simultaneously engineer *rpl36a^P56Q^* and *rtls^PTS1^* alleles and then selected for cycloheximide resistance, sequence heterogeneity was detected at *rtls* exon 4 (indicated by N at positions were the base cannot automatically be assigned) and revealed that the *rtls^PTS1^* allele was present at 16.5% (Population 1) or 10.4% (Population 2) in populations of cells from two independent experiments in which selection for cycloheximide resistant cells was performed after co-editing *rpl36a^P56Q^* and *rtls^PTS1^* alleles. Allele frequency was estimated by unmixing wild-type and *rtls^PTS1^* alleles in electropherograms from Sanger sequencing (Brinkman et al., 2018). Carets indicate site targeted cleavage by *Sp*Cas9. (B) Mutations in *rosetteless* eliminate rosette development. Rosette development in cell populations (N=500 cells for each of three independent replicates) shows that wild-type and *rpl36a^P56Q^* develop into rosettes in the presence of RIFs while *rtls^PTS1^ rpl36a^P56Q^* and *rtls^tl1^* do not. (C-F) Mutations in *rosetteless* prevent the secretion of Rosetteless protein at the basal end of cells and into the interior or rosettes. Immunofluorescent staining for Rosetteless (green), alpha tubulin (gray), and actin (magenta) in wild-type (C), *rpl36a^P56Q^* (D), *rtls^PTS1^ rpl36a^P56Q^* (E), and *rtls^tl1^* (F) strains with (C”-F”) and without (C’-F’) rosette induction. Rosetteless protein localizes in the interior of rosettes (arrow) in wild-type and *rpl36a^P56Q^* but not *rtls^PTS1^ rpl36a^P56Q^* and *rtls^tl1^*.

**Figure S4:**
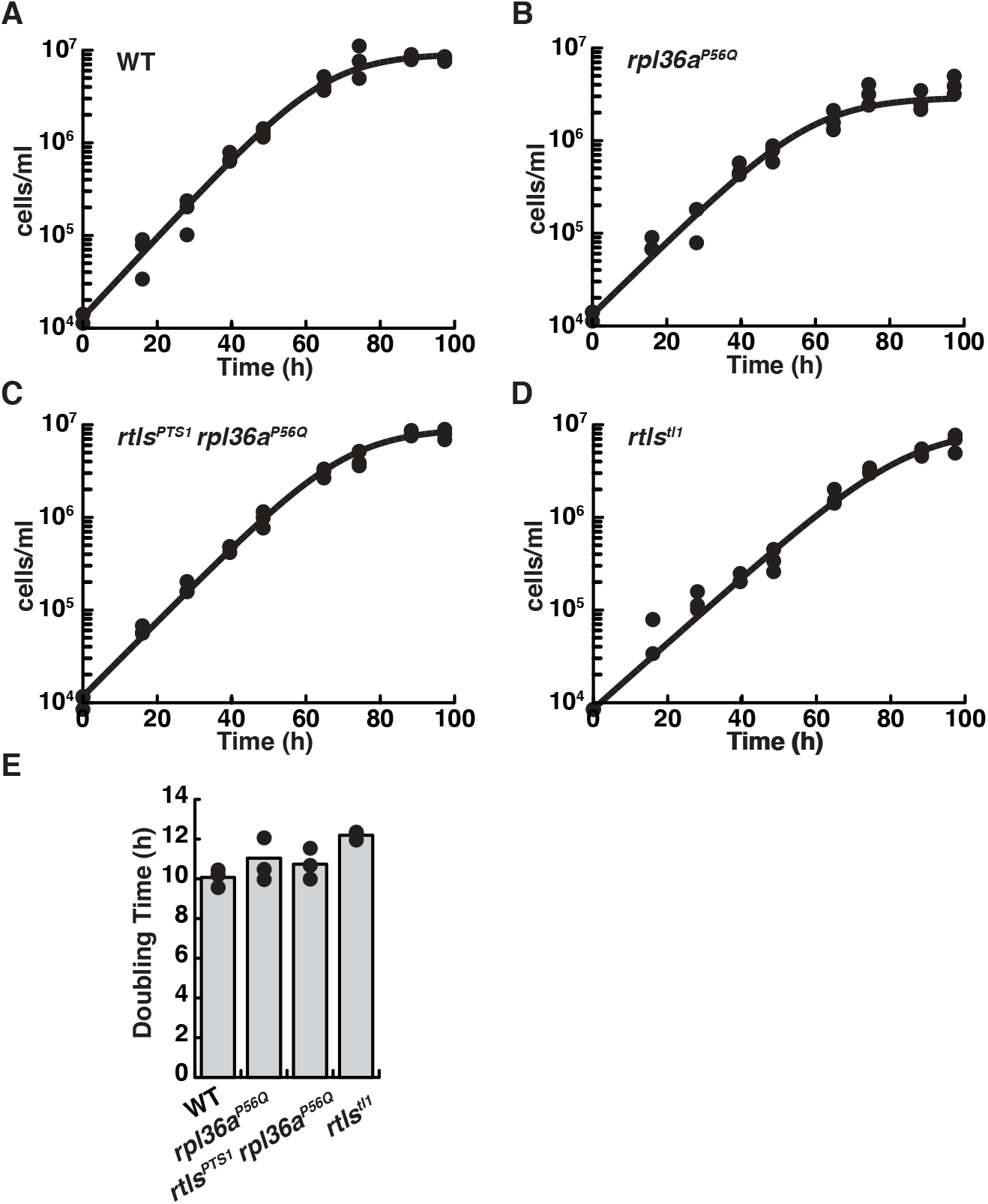
Wild-type and mutant strains proliferate similarly. Growth curves for wild-type (A), *rpl36a^P56Q^* (B), *rtls^PTS1^ rpl36a^P56Q^* (C), and *rtls^tl1^* (D) show similar rates of proliferation. The growth for each strain was characterized by seeding cells at a density of 1×10^4^ cells per ml and determination the cell concentration every ∼12 hours. For each time point, triplicate measures were taken. Each replicate growth trajectory was fit with the logistic equation to calculate the doubling time (E). An analysis of variance (ANOVA) between samples showed no significant differences between growth rates.

**Figure S5:**
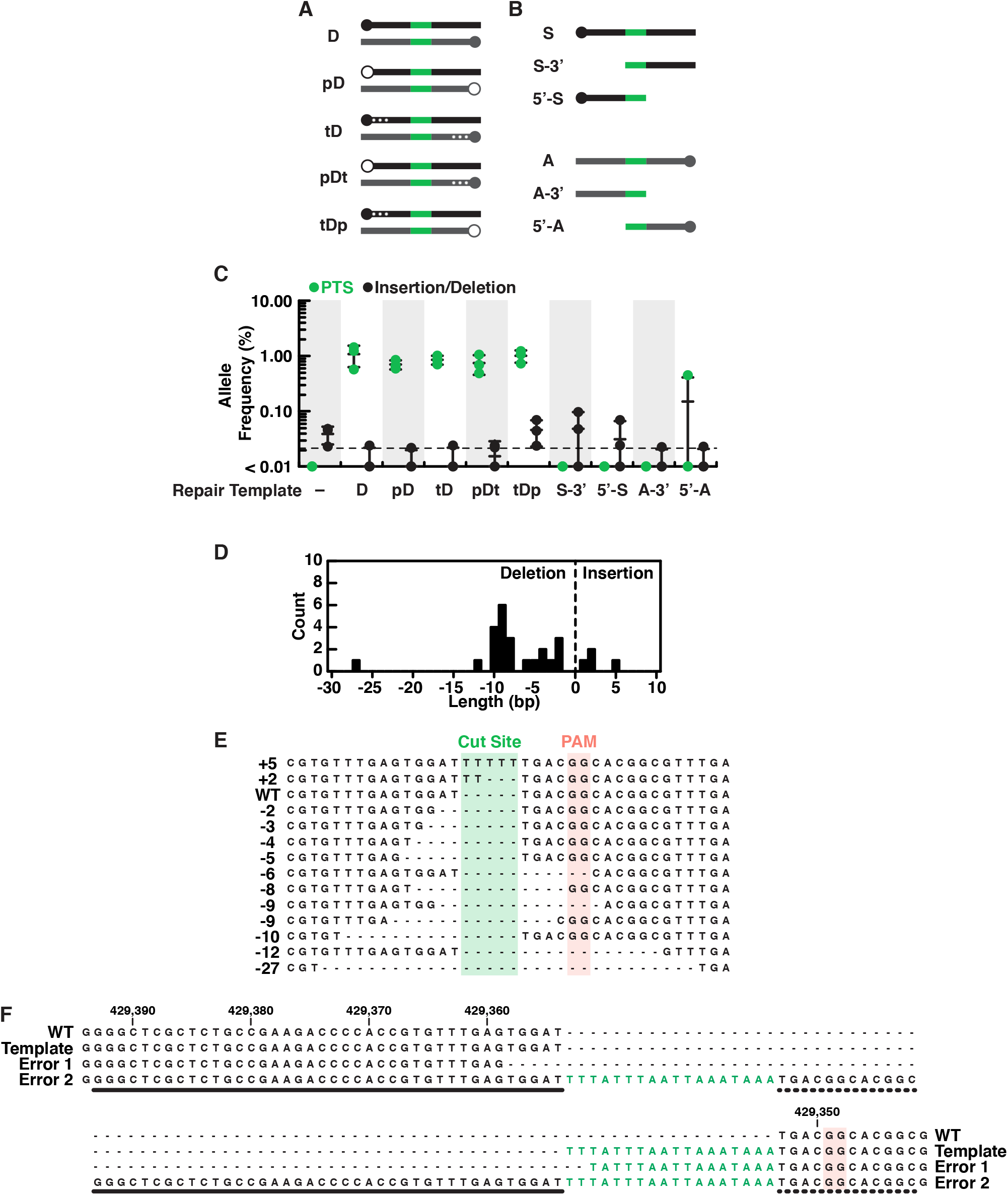
Characterization of editing outcomes at the *rosetteless* locus with different types of repair templates. (A) Double-stranded DNA repair templates (black indicates homology arms from the sense strand, gray indicates homology arms from the antisense strand and green is the PTS as in Fig. 4) were designed with phosphorylated 5’ ends (indicated with open circles at the 5’ end and a ‘p’ in template names; closed circles indicate unphosphorylated ends) or three phosphorothioate bonds between bases at the 5’ end (indicated with asterisks in diagrams and a ‘t’ in template names). We hypothesized that phosphorylated templates would be more susceptible to nucleases and phosphorothioate bonds less susceptible (Renaud et al., 2016; Yu et al., 2019), altering their utility as repair substrates *in vivo*, yet *S. rosetta* used all double stranded templates with similar efficiency (see panel C). (B) We also designed a panel of single-stranded repair templates (colors as in panel A) that lacked 5’ or 3’ arms (Paix et al., 2017) and found that both arms of homology are required for efficient template-mediated genome editing in *S. rosetta* (see panel C). (C) A comparison of DNA repair templates revealed that *S. rosetta* efficiently uses double stranded DNA templates during DNA repair and requires both arms of homology for single-stranded DNA templates. Frequencies of alleles containing either the PTS (green) or insertion/deletion mutations (black) are shown for genome editing experiments based on each of the templates described in panels A and B. Genome editing in the presence of double-stranded DNA templates favored the template-directed DNA repair. The use of phosphorylated double-stranded DNA templates or double-stranded DNA templates templates with phosphorothioate bonds (see panel A) did not increase editing efficiency over unmodified double-stranded DNA templates. We also found that removing 5’ or 3’ homology arms from single stranded templates (see panel B) almost completely eliminated efficient editing as compared to single-stranded templates with both homology arms (Fig. 4C). Each editing experiment was performed three independent times. (D-E) An aggregate analysis of insertion/deletion mutations identified in deep sequencing of genome editing experiments. (D) A histogram shows the length and frequency of insertion and deletion mutations. (E) A sequence alignment or representative insertion and deletion mutations from each size of insertion/deletion mutations. Notably, the most frequent deletions (8-10 bases) occur at dinucleotide repeats, suggesting that microhomologies may promote deletions after double-stranded breaks. (F) An extreme example of templated repair suggests that *S. rosetta* may incorporate larger insertions. One mutation identified in deep sequencing shows an 88-base insertion, with the insertion featuring two PTS sequences with an intervening region that has some homology to sequences to the left (thick line) and right (dotted line) of the double stranded break. Although we are unsure of the mechanism that led to this mutation, its presence suggests that large mutations could be incorporated into *S. rosetta* via genome editing.

### Supplementary Tables and Scripts (see attached .xlsx and .sc files)

**Table S1: Media recipes**

Recipes for making artificial seawater (Hallegraeff et al., 2004; Skelton et al., 2009), high nutrient media (modified from King et al., 2009; Levin and King, 2013; Booth et al., 2018), and low nutrient media.

**Table S2: Oligonucleotide sequences**

Sequences for gRNAs, repair oligonucleotides, and primers that were used to construct and to validate genome edited strains.

**Table S3: *S. rosetta* strains**

Genotypes and sources of *S. rosetta* strains used in this study.

**Table S4: Deep sequencing library primers**

Sequences for primers (adapted from Lin et al., 2014) used to generate libraries for deep sequencing (Fig. 4 and S5)

**Script S1: Quantifying DNA repair outcomes**

BASH script for quantifying the frequency of repair outcomes from deep sequencing data that was preprocessed and aligned in a Galaxy server (Afgan et al., 2018).

## References

Abedin M, King N. 2008. The Premetazoan Ancestry of Cadherins. Science 319:946–948. doi:10.1126/science.1151084

Afgan E, Baker D, Batut B, van den Beek M, Bouvier D, Čech M, Chilton J, Clements D, Coraor N, Grüning BA, Guerler A, Hillman-Jackson J, Hiltemann S, Jalili V, Rasche H, Soranzo N, Goecks J, Taylor J, Nekrutenko A, Blankenberg D. 2018. The Galaxy platform for accessible, reproducible and collaborative biomedical analyses: 2018 update. Nucleic Acids Res 46:W537–W544. doi:10.1093/nar/gky379

Alegado RA, Brown LW, Cao S, Dermenjian RK, Zuzow R, Fairclough SR, Clardy J, King N. 2012. A bacterial sulfonolipid triggers multicellular development in the closest living relatives of animals. eLife 1:e00013. doi:10.7554/eLife.00013

Alegado RA, Grabenstatter JD, Zuzow R, Morris A, Huang SY, Summons RE, King N. 2013. *Algoriphagus machipongonensis* sp. nov., co-isolated with a colonial choanoflagellate. Int J Syst Evol Microbiol 63:163–168. doi:10.1099/ijs.0.038646-0

Amacher JF, Hobbs HT, Cantor AC, Shah L, Rivero M-J, Mulchand SA, Kuriyan J. 2018. Phosphorylation control of the ubiquitin ligase Cbl is conserved in choanoflagellates. Protein Sci 27:923–932. doi:10.1002/pro.3397

Ares M, Bruns PJ. 1978. Isolation and genetic characterization of a mutation affecting ribosomal resistance to cycloheximide in Tetrahymena. Genetics 90:463–474.

Arribere JA, Bell RT, Fu BXH, Artiles KL, Hartman PS, Fire AZ. 2014. Efficient Marker-Free Recovery of Custom Genetic Modifications with CRISPR/Cas9 in *Caenorhabditis elegans*. Genetics 198:837–846. doi:10.1534/genetics.114.169730

Bae J-H, Sung BH, Sohn J-H. 2018. Site saturation mutagenesis of ribosomal protein L42 at 56th residue and application as a consecutive selection marker for cycloheximide resistance in yeast. FEMS Microbiol Lett 365. doi:10.1093/femsle/fny066

Bhattacharyya M, Stratton MM, Going CC, McSpadden ED, Huang Y, Susa AC, Elleman A, Cao YM, Pappireddi N, Burkhardt P, Gee CL, Barros T, Schulman H, Williams ER, Kuriyan J. 2016. Molecular mechanism of activation-triggered subunit exchange in Ca2+/calmodulin-dependent protein kinase II. eLife 5:e13405. doi:10.7554/eLife.13405

Bibikova M, Carroll D, Segal DJ, Trautman JK, Smith J, Kim Y-G, Chandrasegaran S. 2001. Stimulation of Homologous Recombination through Targeted Cleavage by Chimeric Nucleases. Mol Cell Biol 21:289–297. doi:10.1128/MCB.21.1.289-297.2001

Booth DS, Szmidt-Middleton H, King N. 2018. Choanoflagellate transfection illuminates their cell biology and the ancestry of animal septins. Mol Biol Cell mbcE18080514. doi:10.1091/mbc.E18-08-0514

Brunet T, King N. 2017. The Origin of Animal Multicellularity and Cell Differentiation. Dev Cell 43:124–140. doi:10.1016/j.devcel.2017.09.016

Brunet T, Larson BT, Linden TA, Vermeij MJA, McDonald K, King N. 2019. Light-regulated collective contractility in a multicellular choanoflagellate. Science 366:326–334. doi:10.1126/science.aay2346

Burger G, Forget L, Zhu Y, Gray MW, Lang BF. 2003. Unique mitochondrial genome architecture in unicellular relatives of animals. Proc Natl Acad Sci 100:892–897. doi:10.1073/pnas.0336115100

Carr M, Leadbeater BSC, Hassan R, Nelson M, Baldauf SL. 2008. Molecular phylogeny of choanoflagellates, the sister group to Metazoa. Proc Natl Acad Sci 105:16641–16646. doi:10.1073/pnas.0801667105

Chen B, Gilbert LA, Cimini BA, Schnitzbauer J, Zhang W, Li G-W, Park J, Blackburn EH, Weissman JS, Qi LS, Huang B. 2013. Dynamic imaging of genomic loci in living human cells by an optimized CRISPR/Cas system. Cell 155:1479–1491. doi:10.1016/j.cell.2013.12.001

Chomczynski P, Rymaszewski M. 2006. Alkaline polyethylene glycol-based method for direct PCR from bacteria, eukaryotic tissue samples, and whole blood. BioTechniques 40:454, 456, 458. doi:10.2144/000112149

Choulika A, Perrin A, Dujon B, Nicolas JF. 1995. Induction of homologous recombination in mammalian chromosomes by using the I-SceI system of *Saccharomyces cerevisiae*. Mol Cell Biol 15:1968–1973.

Colgren J, Nichols SA. 2019. The significance of sponges for comparative studies of developmental evolution. WIREs Dev Biol n/a:e359. doi:10.1002/wdev.359

Cong L, Ran FA, Cox D, Lin S, Barretto R, Habib N, Hsu PD, Wu X, Jiang W, Marraffini LA, Zhang F. 2013. Multiplex Genome Engineering Using CRISPR/Cas Systems. Science 339:819–823. doi:10.1126/science.1231143

Cummings RD, McEver RP. 2015. C-Type Lectins In: Varki A, Cummings RD, Esko JD, Stanley P, Hart GW, Aebi M, Darvill AG, Kinoshita T, Packer NH, Prestegard JH, Schnaar RL, Seeberger PH, editors. Essentials of Glycobiology. Cold Spring Harbor (NY): Cold Spring Harbor Laboratory Press.

Dayel MJ, Alegado RA, Fairclough SR, Levin TC, Nichols SA, McDonald K, King N. 2011. Cell differentiation and morphogenesis in the colony-forming choanoflagellate *Salpingoeca rosetta*. Dev Biol 357:73–82. doi:10.1016/j.ydbio.2011.06.003

Dehoux P, Davies J, Cannon M. 1993. Natural cycloheximide resistance in yeast. The role of ribosomal protein L41. Eur J Biochem 213:841–848.

Doench JG, Hartenian E, Graham DB, Tothova Z, Hegde M, Smith I, Sullender M, Ebert BL, Xavier RJ, Root DE. 2014. Rational design of highly active sgRNAs for CRISPR-Cas9-mediated gene inactivation. Nat Biotechnol 32:1262–1267. doi:10.1038/nbt.3026

Drickamer K, Fadden AJ. 2002. Genomic analysis of C-type lectins. Biochem Soc Symp 59–72. doi:10.1042/bss0690059

Fairclough SR, Chen Z, Kramer E, Zeng Q, Young S, Robertson HM, Begovic E, Richter DJ, Russ C, Westbrook MJ, Manning G, Lang BF, Haas B, Nusbaum C, King N. 2013. Premetazoan genome evolution and the regulation of cell differentiation in the choanoflagellate *Salpingoeca rosetta*. Genome Biol 14:R15. doi:10.1186/gb-2013-14-2-r15

Fairclough SR, Dayel MJ, King N. 2010. Multicellular development in a choanoflagellate. Curr Biol 20:R875–R876. doi:10.1016/j.cub.2010.09.014

Ferenczi A, Pyott DE, Xipnitou A, Molnar A. 2017. Efficient targeted DNA editing and replacement in Chlamydomonas reinhardtii using Cpf1 ribonucleoproteins and single-stranded DNA. Proc Natl Acad Sci U S A 114:13567–13572. doi:10.1073/pnas.1710597114

Foster AJ, Martin-Urdiroz M, Yan X, Wright HS, Soanes DM, Talbot NJ. 2018. CRISPR-Cas9 ribonucleoprotein-mediated co-editing and counterselection in the rice blast fungus. Sci Rep 8:14355. doi:10.1038/s41598-018-32702-w

Gandhi S, Haeussler M, Razy-Krajka F, Christiaen L, Stolfi A. 2017. Evaluation and rational design of guide RNAs for efficient CRISPR/Cas9-mediated mutagenesis in Ciona. Dev Biol 425:8–20. doi:10.1016/j.ydbio.2017.03.003

Garreau de Loubresse N, Prokhorova I, Holtkamp W, Rodnina MV, Yusupova G, Yusupov M. 2014. Structural basis for the inhibition of the eukaryotic ribosome. Nature 513:517–522. doi:10.1038/nature13737

Gilles AF, Averof M. 2014. Functional genetics for all: engineered nucleases, CRISPR and the gene editing revolution. EvoDevo 5:43. doi:10.1186/2041-9139-5-43

Grau-Bové X, Torruella G, Donachie S, Suga H, Leonard G, Richards TA, Ruiz-Trillo I. 2017. Dynamics of genomic innovation in the unicellular ancestry of animals. eLife 6:e26036. doi:10.7554/eLife.26036

Harrison MM, Jenkins BV, O’Connor-Giles KM, Wildonger J. 2014. A CRISPR view of development. Genes Dev 28:1859–1872. doi:10.1101/gad.248252.114

Hoffmeyer TT, Burkhardt P. 2016. Choanoflagellate models — *Monosiga brevicollis* and *Salpingoeca rosetta*. *Curr Opin Genet Dev*, Developmental mechanisms, patterning and evolution 39:42–47. doi:10.1016/j.gde.2016.05.016

Horlbeck MA, Gilbert LA, Villalta JE, Adamson B, Pak RA, Chen Y, Fields AP, Park CY, Corn JE, Kampmann M, Weissman JS. 2016. Compact and highly active next-generation libraries for CRISPR-mediated gene repression and activation. eLife 5. doi:10.7554/eLife.19760

Jacobs JZ, Ciccaglione KM, Tournier V, Zaratiegui M. 2014. Implementation of the CRISPR-Cas9 system in fission yeast. Nat Commun 5:1–5. doi:10.1038/ncomms6344

Jiang F, Taylor DW, Chen JS, Kornfeld JE, Zhou K, Thompson AJ, Nogales E, Doudna JA. 2016. Structures of a CRISPR-Cas9 R-loop complex primed for DNA cleavage. Science 351:867–871. doi:10.1126/science.aad8282

Jiang W, Brueggeman AJ, Horken KM, Plucinak TM, Weeks DP. 2014. Successful Transient Expression of Cas9 and Single Guide RNA Genes in *Chlamydomonas reinhardtii*. Eukaryot Cell 13:1465–1469. doi:10.1128/EC.00213-14

Jinek M, Chylinski K, Fonfara I, Hauer M, Doudna JA, Charpentier E. 2012. A programmable dual-RNA-guided DNA endonuclease in adaptive bacterial immunity. Science 337:816–821. doi:10.1126/science.1225829

Jinek M, East A, Cheng A, Lin S, Ma E, Doudna J. 2013. RNA-programmed genome editing in human cells. eLife 2:e00471. doi:10.7554/eLife.00471

Kaur K, Gupta AK, Rajput A, Kumar M. 2016. ge-CRISPR - An integrated pipeline for the prediction and analysis of sgRNAs genome editing efficiency for CRISPR/Cas system. Sci Rep 6:30870. doi:10.1038/srep30870

Kawai S, Murao S, Mochizuki M, Shibuya I, Yano K, Takagi M. 1992. Drastic alteration of cycloheximide sensitivity by substitution of one amino acid in the L41 ribosomal protein of yeasts. J Bacteriol 174:254–262.

Kim H, Ishidate T, Ghanta KS, Seth M, Conte D, Shirayama M, Mello CC. 2014. A Co-CRISPR Strategy for Efficient Genome Editing in *Caenorhabditis elegans*. Genetics 197:1069–1080. doi:10.1534/genetics.114.166389

Kim IG, Nam SK, Sohn JH, Rhee SK, An GH, Lee SH, Choi ES. 1998. Cloning of the ribosomal protein L41 gene of *Phaffia rhodozyma* and its use a drug resistance marker for transformation. Appl Environ Microbiol 64:1947–1949.

King N. 2004. The Unicellular Ancestry of Animal Development. Dev Cell 7:313–325. doi:10.1016/j.devcel.2004.08.010

King N, Hittinger CT, Carroll SB. 2003. Evolution of Key Cell Signaling and Adhesion Protein Families Predates Animal Origins. Science 301:361–363. doi:10.1126/science.1083853

King N, Westbrook MJ, Young SL, Kuo A, Abedin M, Chapman J, Fairclough S, Hellsten U, Isogai Y, Letunic I, Marr M, Pincus D, Putnam N, Rokas A, Wright KJ, Zuzow R, Dirks W, Good M, Goodstein D, Lemons D, Li W, Lyons JB, Morris A, Nichols S, Richter DJ, Salamov A, Sequencing JGI, Bork P, Lim WA, Manning G, Miller WT, McGinnis W, Shapiro H, Tjian R, Grigoriev IV, Rokhsar D. 2008. The genome of the choanoflagellate *Monosiga brevicollis* and the origin of metazoans. Nature 451:783–788. doi:10.1038/nature06617

King N, Young SL, Abedin M, Carr M, Leadbeater BSC. 2009. Starting and Maintaining *Monosiga brevicollis* Cultures. Cold Spring Harb Protoc 2009:pdb.prot5148. doi:10.1101/pdb.prot5148

Kondo K, Saito T, Kajiwara S, Takagi M, Misawa N. 1995. A transformation system for the yeast *Candida utilis*: use of a modified endogenous ribosomal protein gene as a drug-resistant marker and ribosomal DNA as an integration target for vector DNA. J Bacteriol 177:7171–7177.

Lang BF, O’Kelly C, Nerad T, Gray MW, Burger G. 2002. The Closest Unicellular Relatives of Animals. Curr Biol 12:1773–1778. doi:10.1016/S0960-9822(02)01187-9

Larson BT, Ruiz-Herrero T, Lee S, Kumar S, Mahadevan L, King N. 2020. Biophysical principles of choanoflagellate self-organization. Proc Natl Acad Sci 117:1303–1311. doi:10.1073/pnas.1909447117

Laundon D, Larson BT, McDonald K, King N, Burkhardt P. 2019. The architecture of cell differentiation in choanoflagellates and sponge choanocytes. PLOS Biol 17:e3000226. doi:10.1371/journal.pbio.3000226

Leadbeater BSC. 2015. The choanoflagellates: evolution, biology, and ecology. Cambridge, United Kingdom: Cambridge University Press.

Levin TC, Greaney AJ, Wetzel L, King N. 2014. The Rosetteless gene controls development in the choanoflagellate *S. rosetta*. eLife 3. doi:10.7554/eLife.04070

Levin TC, King N. 2013. Evidence for Sex and Recombination in the Choanoflagellate *Salpingoeca rosetta*. Curr Biol 23:2176–2180. doi:10.1016/j.cub.2013.08.061

Li H, Beckman KA, Pessino V, Huang B, Weissman JS, Leonetti MD. 2019. Design and specificity of long ssDNA donors for CRISPR-based knock-in. bioRxiv 178905. doi:10.1101/178905

Lin S, Staahl BT, Alla RK, Doudna JA. 2014. Enhanced homology-directed human genome engineering by controlled timing of CRISPR/Cas9 delivery. eLife 3:e04766. doi:10.7554/eLife.04766

Liu X, Homma A, Sayadi J, Yang S, Ohashi J, Takumi T. 2016. Sequence features associated with the cleavage efficiency of CRISPR/Cas9 system. Sci Rep 6:19675. doi:10.1038/srep19675

Lorenz R, Bernhart SH, Höner Zu Siederdissen C, Tafer H, Flamm C, Stadler PF, Hofacker IL. 2011. ViennaRNA Package 2.0. Algorithms Mol Biol AMB 6:26. doi:10.1186/1748-7188-6-26

Manning G, Young SL, Miller WT, Zhai Y. 2008. The protist, Monosiga brevicollis has a tyrosine kinase signaling network more elaborate and diverse than found in any known metazoan. Proc Natl Acad Sci 105:9674. doi:10.1073/pnas.0801314105

Marron AO, Alston MJ, Heavens D, Akam M, Caccamo M, Holland PWH, Walker G. 2013. A family of diatom-like silicon transporters in the siliceous loricate choanoflagellates. Proc R Soc B Biol Sci 280:20122543. doi:10.1098/rspb.2012.2543

Mendoza A de, Sebé-Pedrós A, Šestak MS, Matejčić M, Torruella G, Domazet-Lošo T, Ruiz-Trillo I. 2013. Transcription factor evolution in eukaryotes and the assembly of the regulatory toolkit in multicellular lineages. Proc Natl Acad Sci 110:E4858–E4866. doi:10.1073/pnas.1311818110

Momose T, Concordet J-P. 2016. Diving into marine genomics with CRISPR/Cas9 systems. Mar Genomics 30:55–65. doi:10.1016/j.margen.2016.10.003

Moreno-Mateos MA, Vejnar CE, Beaudoin J-D, Fernandez JP, Mis EK, Khokha MK, Giraldez AJ. 2015. CRISPRscan: designing highly efficient sgRNAs for CRISPR-Cas9 targeting in vivo. Nat Methods 12:982–988. doi:10.1038/nmeth.3543

Nichols SA, Roberts BW, Richter DJ, Fairclough SR, King N. 2012. Origin of metazoan cadherin diversity and the antiquity of the classical cadherin/β-catenin complex. Proc Natl Acad Sci 109:13046–13051. doi:10.1073/pnas.1120685109

Oberacker P, Stepper P, Bond DM, Höhn S, Focken J, Meyer V, Schelle L, Sugrue VJ, Jeunen G-J, Moser T, Hore SR, von Meyenn F, Hipp K, Hore TA, Jurkowski TP. 2019. Bio-On-Magnetic-Beads (BOMB): Open platform for high-throughput nucleic acid extraction and manipulation. PLoS Biol 17:e3000107. doi:10.1371/journal.pbio.3000107

Okamoto S, Amaishi Y, Maki I, Enoki T, Mineno J. 2019. Highly efficient genome editing for single-base substitutions using optimized ssODNs with Cas9-RNPs. Sci Rep 9:1–11. doi:10.1038/s41598-019-41121-4

Paix A, Folkmann A, Goldman DH, Kulaga H, Grzelak MJ, Rasoloson D, Paidemarry S, Green R, Reed RR, Seydoux G. 2017. Precision genome editing using synthesis-dependent repair of Cas9-induced DNA breaks. Proc Natl Acad Sci 114:E10745–E10754. doi:10.1073/pnas.1711979114

Parfrey LW, Lahr DJG, Knoll AH, Katz LA. 2011. Estimating the timing of early eukaryotic diversification with multigene molecular clocks. Proc Natl Acad Sci U S A 108:13624–13629. doi:10.1073/pnas.1110633108

Peña JF, Alié A, Richter DJ, Wang L, Funayama N, Nichols SA. 2016. Conserved expression of vertebrate microvillar gene homologs in choanocytes of freshwater sponges. EvoDevo 7:13. doi:10.1186/s13227-016-0050-x

Peng D, Tarleton R. 2015. EuPaGDT: a web tool tailored to design CRISPR guide RNAs for eukaryotic pathogens. Microb Genomics 1:e000033. doi:10.1099/mgen.0.000033

Pincus D, Letunic I, Bork P, Lim WA. 2008. Evolution of the phospho-tyrosine signaling machinery in premetazoan lineages. Proc Natl Acad Sci 105:9680. doi:10.1073/pnas.0803161105

Renaud J-B, Boix C, Charpentier M, De Cian A, Cochennec J, Duvernois-Berthet E, Perrouault L, Tesson L, Edouard J, Thinard R, Cherifi Y, Menoret S, Fontanière S, de Crozé N, Fraichard A, Sohm F, Anegon I, Concordet J-P, Giovannangeli C. 2016. Improved Genome Editing Efficiency and Flexibility Using Modified Oligonucleotides with TALEN and CRISPR-Cas9 Nucleases. Cell Rep 14:2263–2272. doi:10.1016/j.celrep.2016.02.018

Richardson CD, Ray GJ, DeWitt MA, Curie GL, Corn JE. 2016. Enhancing homology-directed genome editing by catalytically active and inactive CRISPR-Cas9 using asymmetric donor DNA. Nat Biotechnol 34:339–344. doi:10.1038/nbt.3481

Richter DJ, Fozouni P, Eisen MB, King N. 2018. Gene family innovation, conservation and loss on the animal stem lineage. eLife 7:e34226. doi:10.7554/eLife.34226

Rouet P, Smih F, Jasin M. 1994. Introduction of double-strand breaks into the genome of mouse cells by expression of a rare-cutting endonuclease. Mol Cell Biol 14:8096–8106. doi:10.1128/MCB.14.12.8096

Ruiz-Trillo I, Roger AJ, Burger G, Gray MW, Lang BF. 2008. A Phylogenomic Investigation into the Origin of Metazoa. Mol Biol Evol 25:664–672. doi:10.1093/molbev/msn006

Schindelin J, Arganda-Carreras I, Frise E, Kaynig V, Longair M, Pietzsch T, Preibisch S, Rueden C, Saalfeld S, Schmid B, Tinevez J-Y, White DJ, Hartenstein V, Eliceiri K, Tomancak P, Cardona A. 2012. Fiji: an open-source platform for biological-image analysis. Nat Methods 9:676–682. doi:10.1038/nmeth.2019

Sebé-Pedrós A, Burkhardt P, Sánchez-Pons N, Fairclough SR, Lang BF, King N, Ruiz-Trillo I. 2013. Insights into the Origin of Metazoan Filopodia and Microvilli. Mol Biol Evol 30:2013–2023. doi:10.1093/molbev/mst110

Sebé-Pedrós A, Degnan BM, Ruiz-Trillo I. 2017. The origin of Metazoa: a unicellular perspective. Nat Rev Genet 18:498–512. doi:10.1038/nrg.2017.21

Shin S-E, Lim J-M, Koh HG, Kim EK, Kang NK, Jeon S, Kwon S, Shin W-S, Lee B, Hwangbo K, Kim J, Ye SH, Yun J-Y, Seo H, Oh H-M, Kim K-J, Kim J-S, Jeong W-J, Chang YK, Jeong B-R. 2016. CRISPR/Cas9-induced knockout and knock-in mutations in *Chlamydomonas reinhardtii*. Sci Rep 6:27810. doi:10.1038/srep27810

Stevens DR, Atteia A, Franzén LG, Purton S. 2001. Cycloheximide resistance conferred by novel mutations in ribosomal protein L41 of *Chlamydomonas reinhardtii*. Mol Gen Genet MGG 264:790–795.

Sutton CA, Ares M, Hallberg RL. 1978. Cycloheximide resistance can be mediated through either ribosomal subunit. Proc Natl Acad Sci U S A 75:3158–3162.

Thyme SB, Akhmetova L, Montague TG, Valen E, Schier AF. 2016. Internal guide RNA interactions interfere with Cas9-mediated cleavage. Nat Commun 7:11750. doi:10.1038/ncomms11750

Ward JD. 2015. Rapid and precise engineering of the *Caenorhabditis elegans* genome with lethal mutation co-conversion and inactivation of NHEJ repair. Genetics 199:363–377. doi:10.1534/genetics.114.172361

Wetzel LA, Levin TC, Hulett RE, Chan D, King GA, Aldayafleh R, Booth DS, Sigg MA, King N. 2018. Predicted glycosyltransferases promote development and prevent spurious cell clumping in the choanoflagellate *S. rosetta*. eLife 7. doi:10.7554/eLife.41482

Woznica A, Cantley AM, Beemelmanns C, Freinkman E, Clardy J, King N. 2016. Bacterial lipids activate, synergize, and inhibit a developmental switch in choanoflagellates. Proc Natl Acad Sci U S A 113:7894–7899. doi:10.1073/pnas.1605015113

Woznica A, Gerdt JP, Hulett RE, Clardy J, King N. 2017. Mating in the Closest Living Relatives of Animals Is Induced by a Bacterial Chondroitinase. Cell 170:1175–1183.e11. doi:10.1016/j.cell.2017.08.005

Wu X, Scott DA, Kriz AJ, Chiu AC, Hsu PD, Dadon DB, Cheng AW, Trevino AE, Konermann S, Chen S, Jaenisch R, Zhang F, Sharp PA. 2014. Genome-wide binding of the CRISPR endonuclease Cas9 in mammalian cells. Nat Biotechnol 32:670–676. doi:10.1038/nbt.2889

Xu H, Xiao T, Chen C-H, Li W, Meyer CA, Wu Q, Wu D, Cong L, Zhang F, Liu JS, Brown M, Liu XS. 2015. Sequence determinants of improved CRISPR sgRNA design. Genome Res 25:1147–1157. doi:10.1101/gr.191452.115

Yeh CD, Richardson CD, Corn JE. 2019. Advances in genome editing through control of DNA repair pathways. Nat Cell Biol 21:1468–1478. doi:10.1038/s41556-019-0425-z

Young SL, Diolaiti D, Conacci-Sorrell M, Ruiz-Trillo I, Eisenman RN, King N. 2011. Premetazoan Ancestry of the Myc–Max Network. Mol Biol Evol 28:2961–2971. doi:10.1093/molbev/msr132

Yu Y, Guo Y, Tian Q, Lan Y, Yeh H, Zhang M, Tasan I, Jain S, Zhao H. 2019. An efficient gene knock-in strategy using 5′-modified double-stranded DNA donors with short homology arms. Nat Chem Biol 1–4. doi:10.1038/s41589-019-0432-1

## Supplementary References

Atlas RM. 2010. Handbook of Microbiological Media, Fourth Edition. Boca Raton: CRC Press.

Brinkman EK, Kousholt AN, Harmsen T, Leemans C, Chen T, Jonkers J, van Steensel B. 2018. Easy quantification of template-directed CRISPR/Cas9 editing. Nucleic Acids Res 46:e58–e58. doi:10.1093/nar/gky164

Gibson DG, Young L, Chuang R-Y, Venter JC, Hutchison CA, Smith HO. 2009. Enzymatic assembly of DNA molecules up to several hundred kilobases. Nat Methods 6:343–345. doi:10.1038/nmeth.1318

Goddard TD, Huang CC, Ferrin TE. 2005. Software Extensions to UCSF Chimera for Interactive Visualization of Large Molecular Assemblies. Structure 13:473–482. doi:10.1016/j.str.2005.01.006

Hallegraeff GM, Anderson DM, Cembella AD, Enevoldsen HO. 2004. Manual on harmful marine microalgae. Paris: UNESCO.

Milligan JF, Uhlenbeck OC. 1989. Synthesis of small RNAs using T7 RNA polymerase. Methods Enzymol 180:51–62. doi:10.1016/0076-6879(89)80091-6

Sievers F, Wilm A, Dineen D, Gibson TJ, Karplus K, Li W, Lopez R, McWilliam H, Remmert M, Söding J, Thompson JD, Higgins DG. 2011. Fast, scalable generation of high-quality protein multiple sequence alignments using Clustal Omega. Mol Syst Biol 7:539. doi:10.1038/msb.2011.75

Skelton HM, Burkholder JM, Parrow MW. 2009. Axenic Culture of the Heterotrophic Dinoflagellate *Pfiesteria shumwayae* in a Semi-Defined Medium. J Eukaryot Microbiol 56:73–82. doi:10.1111/j.1550-7408.2008.00368.x

Staahl BT, Benekareddy M, Coulon-Bainier C, Banfal AA, Floor SN, Sabo JK, Urnes C, Munares GA, Ghosh A, Doudna JA. 2017. Efficient genome editing in the mouse brain by local delivery of engineered Cas9 ribonucleoprotein complexes. Nat Biotechnol 35:431–434. doi:10.1038/nbt.3806

